# Direct comparison of clathrin-mediated endocytosis in budding and fission yeast reveals conserved and evolvable features

**DOI:** 10.1101/733543

**Authors:** Yidi Sun, Johnannes Schoeneberg, Shirley Chen, Tommy Jiang, Charlotte Kaplan, Ke Xu, Thomas D. Pollard, David G. Drubin

## Abstract

Conserved proteins drive clathrin-mediated endocytosis (CME), which universally involves a burst of actin assembly. To gain fundamental mechanistic insights into this process, a side-by-side quantitative comparison of CME was performed on two distantly related yeast species. Though endocytic protein abundance in *S. pombe* and *S. cerevisiae* are more similar than previously thought, membrane invagination speed and depth are two-fold greater in fission yeast than in budding yeast. In both yeasts, Arp2/3 complex activation drives membrane invagination when triggered by the accumulation of ∼70 WASP molecules. In contrast to budding yeast, WASP-mediated actin nucleation activity plays an essential role in fission yeast endocytosis. Genetics and live-cell imaging revealed core CME spatiodynamic similarities between the two yeasts, though two-zone actin assembly is a fission yeast-specific mechanism, which is not essential for CME. These studies identified conserved CME mechanisms and species-specific adaptations and have broad implications that extend from yeast to humans.

## Introduction

Clathrin-mediated endocytosis (CME) takes up nutrients, regulates responses to extracellular stimuli, and controls the chemical composition and surface area of the plasma membrane. From yeast to humans, several dozen highly conserved proteins are recruited to CME sites in a regular, predictable order to facilitate plasma membrane internalization (Engqvist-Goldstein and Drubin, 2003; Haucke and Kozlov, 2018; Kaksonen and Roux, 2018; McMahon and Boucrot, 2011; Mettlen et al., 2018; Lu et al., 2016).

During CME, cells deform the plasma membrane locally and subsequently pinch off a vesicle into the cytoplasm. But the forces required to achieve membrane deformation and the sizes of the resulting vesicles can vary widely between species and cell types (Lacy et al., 2018; McMahon and Boucrot, 2011). For animal cells, different cell types, and even different areas of the same cell, may exhibit differences in plasma membrane tension (Boulant et al., 2011), affecting the forces required to create a membrane invagination. Both budding and fission yeast have high internal turgor pressure, which strongly opposes plasma membrane invagination necessary for endocytosis. Actin filament assembly mediated by the Arp2/3 (actin-related protein 2/3) complex generates force to overcome plasma membrane tension in animal cells (Boulant et al., 2011; Merrifield et al., 2004; Yarar et al., 2005) and turgor pressure in yeast (Aghamohammadzadeh and Ayscough, 2009; Basu et al., 2014; Kaksonen et al., 2003; Sirotkin et al., 2005). How various organisms utilize a similar set of proteins to satisfy their particular force generation requirements for endocytosis is an unanswered, fundamental question.

Cell biology approaches, combined with the superb genetics of yeast, have advanced our understanding to CME. Electron microscopy (EM) (Buser and Drubin, 2013; Idrissi et al., 2012; Kukulski et al., 2012; Mulholland et al., 1994), super resolution microscopy (Mund et al., 2018) and comprehensive, high frame rate (Picco et al., 2015), live-cell imaging of many of the proteins tagged with fluorescent proteins (Galletta et al., 2008; Kaksonen et al., 2003; Kaksonen et al., 2005; Newpher et al., 2005; Sun et al., 2006) produced a detailed protein recruitment time course that was mapped onto precise morphological stages of membrane deformation (such as invagination length and vesicle release) in budding yeast (Boettner et al., 2011; Goode et al., 2015; Lu et al., 2016). Such information is not yet available for fission yeast. Thus, the size of the CME membrane invagination is not clear in fission yeast. Quantitative fluorescence microscopy of fission yeast provided counts of endocytic proteins over time and described their recruitment kinetics at endocytic sites (Sirotkin, 2010; Arasada, 2011; Berro, 2014; Basu and Chang, 2011). Live-cell super resolution analysis of endocytosis was also performed in fission yeast (Arasada, 2018). While much of what is known about CME was learned from decades of classic cell biology studies in mammalian cells (Schmid, 2014), more recent advanced experimental approaches, such as genome editing, stem cell technology, and sophisticated new imaging modalities, are beginning to make studies similar to those performed in yeast accessible for mammalian cells (Cocucci et al., 2014; Dambournet et al., 2018; Doyon et al., 2011; Grassart et al., 2014; Kadlecova et al., 2017; Liu et al., 2018; Sochacki et al., 2017; Taylor et al., 2011).

The ancestors of budding yeast and fission yeast separated from each other about 330-420 million years ago, which means that these two yeasts are almost as distant from each other as either is to animal cells (Hedges, 2002; Sipiczki, 2000). That actin assembly accompanies CME from yeast to humans highlights an ancient association between actin and CME. Given that relating high-resolution CME protein number and dynamics to membrane morphology is still difficult in mammalian cells, a direct, quantitative comparison of CME in the two yeasts provides a promising approach to reveal the fundamental and evolutionarily adaptable aspects of the process.

Two distinct models for how actin polymerization produces force for endocytosis, a push and pull model for budding yeast (Sun et al., 2006) and a two-zone model for fission yeast (Arasada and Pollard, 2011) have been developed based on several types of quantitative analyses (Figure 1-figure supplement 1) (Lacy et al., 2018). Precise knowledge of endocytic protein numbers and dynamics is essential for understanding basic mechanisms and how they are adapted by evolution. Therefore, direct comparisons of protein numbers and dynamics between the two yeasts using the same methodology promises to provide evolutionary insights into the process and how it can be modified. Quantitation of protein numbers at endocytic sites was a critical, pioneering development in fission yeast studies (Sirotkin et al., 2010). Comparatively lower numbers of homologous proteins were reported present at budding yeast endocytic sites (Picco et al., 2015), and in some cases discrepancies were reported even for the same protein in the same yeast species (Figure 1-table supplement 1) (Basu and Chang, 2011; Chen and Pollard, 2013; Epstein et al., 2018; Galletta et al., 2012; Picco et al., 2015; Sirotkin et al., 2010), highlighting the importance of direct comparisons. Furthermore, the dynamics of some of the proteins including WASp, one of the main nucleation promoting factors (NPFs) for the Arp2/3 complex, were reported to be distinctly different in budding and fission yeast (Kaksonen et al., 2003; Sirotkin et al., 2010).

Here we compared CME in budding and fission yeast side-by-side by recording the dynamics of endocytic protein homologs simultaneously at high frame rates by wide-field fluorescence microscopy. The live-cell imaging data were processed using a centroid tracking method (Sbalzarini and Koumoutsakos, 2005) and then were analyzed quantitatively using newly-developed custom software, which detects the different stages of fission yeast endocytic internalization, such as invagination elongation, scission, and vesicle release. Additionally, the relative abundance of multiple key endocytic proteins in the actin patches of the two yeasts was directly compared and a new extracellular standard particle with 120 copies of GFP (Hsia et al., 2016) was used to calculate the absolute numbers of these proteins at endocytic sites. Three-dimensional stochastic optical reconstruction microscopy (3D-STORM) of fixed fission yeast cells revealed new structural details of endocytic vesicle formation. Finally, we applied two-color live-cell imaging to fission yeast mutants with deletions of the Arp2/3 complex-activating CA domains from WASp or myosin-I to examine the mechanism of endocytic actin assembly.

Our quantitative, direct comparison of budding and fission yeast revealed dramatic, previously unappreciated differences, as well as similarities in CME in these two evolutionarily distant yeasts. These results provide a framework for developing a deeper understanding of CME in all species.

## Results

### Side-by-side quantitative ratiometric imaging reveals modest differences in numbers of homologous proteins at endocytic sites in budding and fission yeast

To accurately compare the numbers of homologous proteins that are recruited to endocytic sites in budding and fission yeast, we first recorded the dynamics of endogenously-tagged fluorescent endocytic proteins in both yeasts simultaneously in the same microscope field. We used fission yeast media to culture both yeasts because fission yeast grow poorly in budding yeast media (data not shown), while budding yeast cells grow robustly in fission yeast media and their endocytic proteins with endogenous fluorescent protein tags behave normally (Figure 1-figure supplement 2). After accurate ratios were determined, we used a new calibration standard to convert the measured fluorescence intensities to protein numbers.

The fluorescence intensities of homologous proteins at endocytic sites over time were quantitatively analyzed in both yeasts (Figure 1-figure supplement 3 for detailed method). In Figure 1, the maximum intensity of homologous proteins at endocytic sites in budding and fission yeast is plotted for seven protein pairs including coat proteins and the endocytic actin machinery. spPan1 and scPan1 are homologues of mammalian intersectins and couple actin assembly to endocytic sites (Bradford et al., 2015; Sun et al., 2015). spEnd4 and scSla2 are homologs of mammalian Hip1R and couple the endocytic actin network to the plasma membrane and clathrin coat (Kaksonen et al., 2003; Sun et al., 2005). The maximum amount of spPan1 is approximately 1.7 times that of scPan1, while spEnd4 and scSla2 are present at endocytic sites in similar numbers (Figure 1A and 1B). spWsp1/scLas17 and spVrp1/scVrp1 are yeast homologs of mammalian WASp and WIP, respectively. The maximum levels of spWsp1 and spVrp1 are modestly higher (1.4 and 1.2 fold, respectively) at endocytic sites than scLas17 and scVrp1 (Figure 1C and 1D). scMyo3 and scMyo5 are budding yeast type I myosins. The combined number of scMyo3 and scMyo5 molecules is ∼1.8 times the number of spMyo1 molecules at endocytic sites (Figure 1E). spArc5 and scArc15 are homologues of mammalian ARPC5, one of the seven subunits of Arp2/3 complex. The numbers of spArc5 and scArc15 molecules recruited to endocytic sites are indistinguishable (Figure 1F). The actin filament crosslinking protein spFim1 is ∼1.2 fold more abundant in fission yeast than the homologous budding yeast protein, scSac6 (Figure 1G).

**Figure 1.**
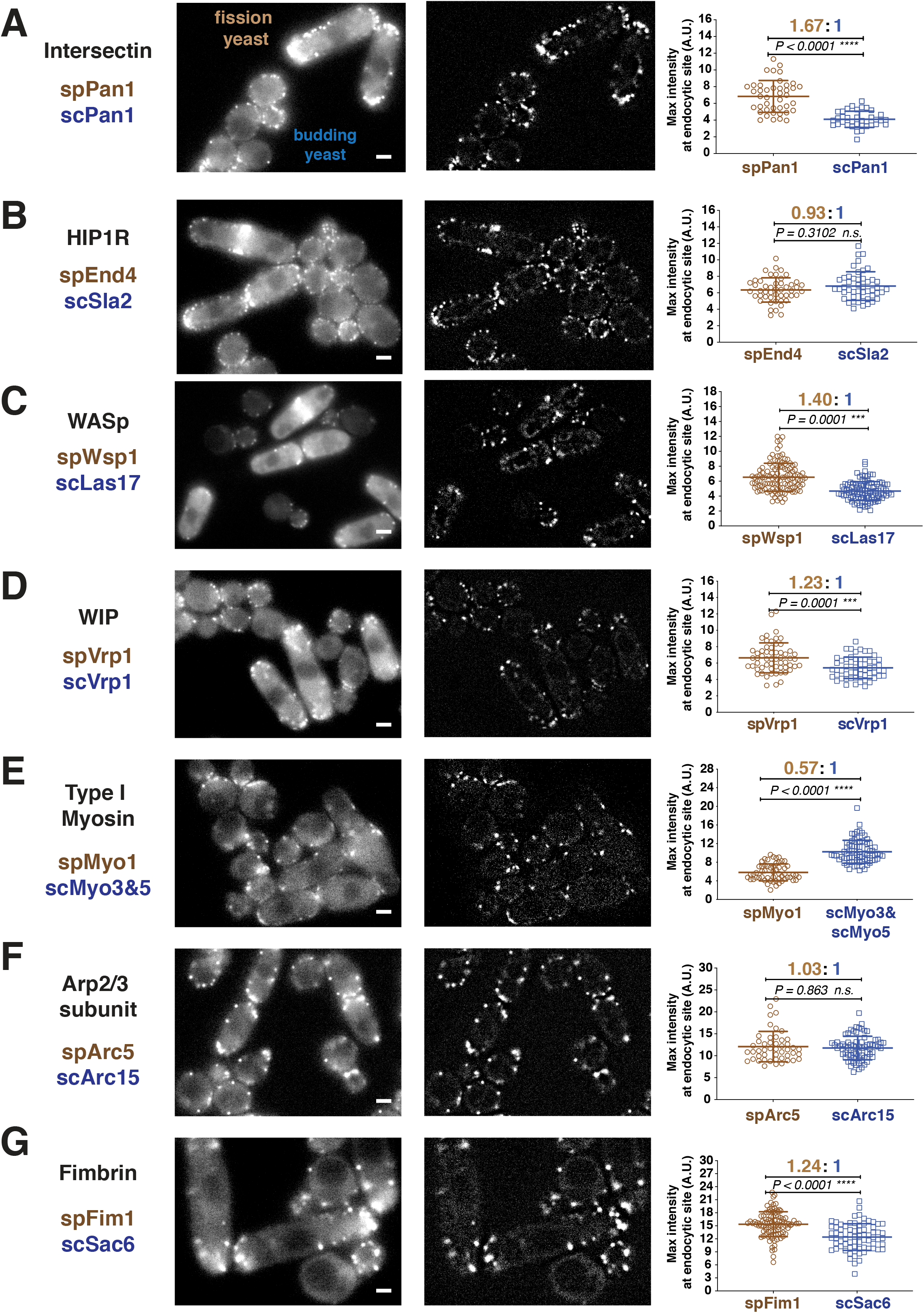
Quantitative side-by-side comparison of levels of homologous proteins at endocytic sites in budding and fission yeast. Single frames from unprocessed (left panel) and processed (middle panel) movies for particle intensity quantification, and average maximum intensity of indicated proteins at endocytic sites (right panel). A-G, Indicated endocytic protein homologues were tagged with GFP or mEGFP in budding yeast or fission yeast, respectively, at the endogenous loci. The maximum intensity for each tagged protein was determined from particle tracking data (see Figure 1-figure supplement 3 for details). mEGFP signal is 14% brighter than GFP (Coffman et al., 2011). The brightness difference was corrected throughout the data analysis. For each indicated protein, at least 50 endocytic sites were examined. n.s. stands for “not significant”. Scale bars on the cell images are 2µm.

These measurements show that the numbers of most of the proteins at endocytic sites are ∼1.2 to ∼1.7 fold higher in fission yeast than their homologues in budding yeast (Figure 1). The exception is type I myosin, which is present at ∼1.8 fold higher levels at budding yeast endocytic sites than at fission yeast sites (Figure 1E). Importantly, this side-by-side comparison (Figure 1 and Figure 2A green bar) shows that the differences in numbers is smaller than reported previously (Picco et al., 2015; Sirotkin et al., 2010) (Figure 2A, grey bar). These results motivated us to reexamine the absolute numbers of the endocytic proteins through direct comparison to a common molecular standard.

**Figure 2.**
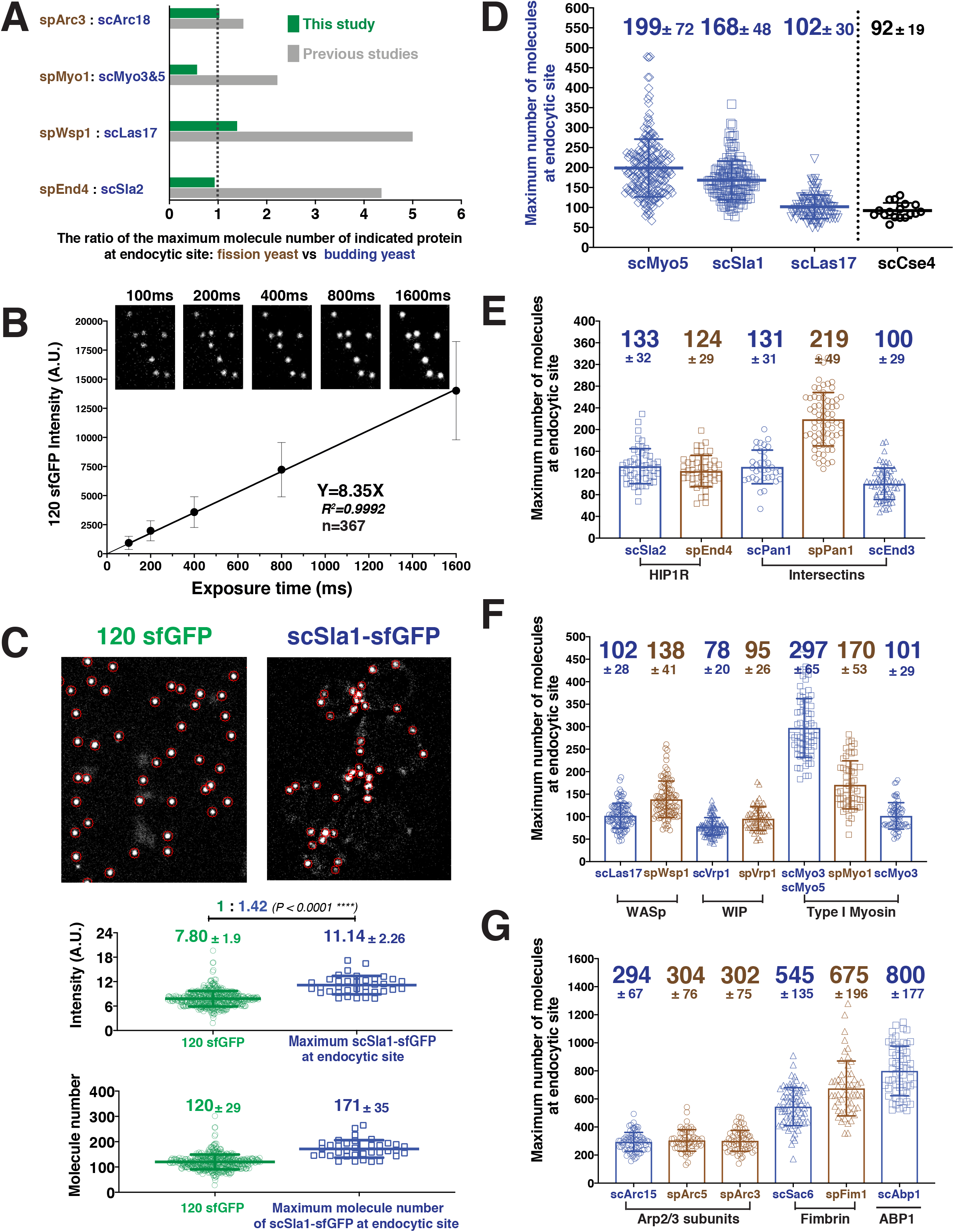
Determining the maximum number of fluorescently-tagged endocytic proteins at endocytic sties by ratiometric comparison of fluorescence intensities to the intensity of 120-sfGFP-tagged nanocages. A, Comparison of ratios of maximum protein levels at endocytic sites for the two yeasts in this study (green) vs in previous studies (grey). B, The mean fluorescence intensity of 120-sfGFP tagged nanocages is linearly proportional to exposure time. Image obtained for indicated exposure times are presented at the top of the graph. C, Ratiometric comparison between the fluorescence intensity of 120-sfGFP tagged nanocages (n=305) and maximum intensity of scSla1-sfGFP at endocytic sites (n=58). Left panel, image of 120-sfGFP tagged nanocages. Right panel, single frame from a processed movie of *scSLA1-sfGFP* cells. The nanocage image and the *scSLA1-sfGFP* movie has been imaged using the same imaging system and the images were processed and analyzed using the same particle tracking parameters. Red circles indicate the sites automatically detected by the tracking program. The intensity (middle) and molecular numbers (bottom) were determined and plotted. D, Maximum molecular number of scMyo5 (n=210), scSla1 (n=152) and scLas17 (n=112) at endocytic sites as well as the molecular number of Cse4 (n=17) on the kinetochore clusters. The molecular numbers were calculated by the ratiometric fluorescence intensity comparison of the indicated sfGFP-tagged proteins and 120-sfGFP tagged nanocages. E-G, Maximum molecular numbers for indicated proteins at endocytic sites in budding (blue) and fission (brown) yeast. The molecular numbers were calculated by the ratiometric fluorescence intensity comparisons using Sla1, Las17, or Myo5 as standards (Figure 2, Figure 2-figure supplement 2). For each indicated protein, at least 50 endocytic sites were examined. The scale bars on the images are 2µm.

### Absolute numbers of proteins at endocytic sites quantified using ratiometric comparison to fluorescence intensity of a 120-sfGFP-tagged nanocage

We calibrated our microscope using as a standard particle a hyperstable, water-soluble 60-subuit protein nanocage (∼25 nm diameter) with in-frame fusions of sfGFP (super-folder Green Fluorescent Protein (GFP)) to both termini of each subunit, so each particle contains 120 copies of sfGFP (Hsia et al., 2016). These particles were expressed in and released from *E. coli* (see Experimental Procedures for details). The distributions of intensities of the 120-sfGFP-tagged nanocages prepared on different days fit Gaussian distributions with similar means and SDs (Figure 2-figure supplement 1). The mean fluorescence intensity was linearly proportional to exposure time over a range from 100 ms to at least 1600 ms per frame (Figure 2B). We used 100-500ms exposure times to image live cells and count molecules. Based on the analysis above, the 120-sfGFP tagged nanocages are expected to enable faithful determination of numbers of fluorescently-tagged proteins present within several fold of 120 using our imaging setup.

Under identical imaging conditions, the mean maximum fluorescence intensities of the endocytic coat proteins Sla1-sfGFP, Myo5-sfGFP and Las17sfGFP at numerous endocytic sites in budding yeast cells, and of hundreds of 120-sfGFP tagged nanocages, were determined (Figure 2C and 2D). Ratiometric comparisons of fluorescence intensity indicate that the mean maximum number of Sla1, Myo5 and Las17 molecules at endocytic sites is 168±48, 199±72, 102±30, respectively (Figure 2D). In budding yeast, the centromere-specific histone H3 variant, Cse4, is a well-accepted standard for counting molecules in live cells (Coffman et al., 2011; Lawrimore et al., 2011). To validate our use of extracellular 120-sfGFP tagged nanocages as a standard for measuring intracellular protein numbers, we determined that the 16-kinetochore clusters present in living anaphase yeast cells contain 92±19 Cse4-sfGFP molecules (Figure 2D), similar to ∼80 to ∼120 molecules counted previously using other calibration methods (Galletta et al., 2012; Lawrimore et al., 2011).

We next determined maximum numbers of various coat proteins (Figure 2E) and other key components at endocytic sites (Figure 2F and 2G) in both yeasts by determining the ratios of fluorescence intensities of these proteins to those of Sla1, Las17, or Myo5. We examined more than 20 pairs of fluorescently-tagged proteins side-by-side (Figure 1, Figure 2-figure supplement 2, and data not shown), yielding absolute numbers for 19 proteins (Figure 2E, 2F, and 2G). To evaluate the quality of these measurements, we used both Sla1-GFP and Las17-GFP as standards to determine the number of Myo3-GFPs present at endocytic sites (Figure 2-figure supplement 3). The numbers determined using both standards were very similar in two independent experiments (Figure 2-figure supplement 3).

Strikingly, the maximum numbers of endocytic proteins we estimated for both budding and fission yeast are generally, to varying degrees, different from those determined in two previous studies (Picco et al., 2015; Sirotkin et al., 2010)(Figure 1-table supplement 1). The numbers we determined for budding yeast were generally 1.5-3 times higher than previously reported (Figure 2E, 2F and 2G) (Picco et al., 2015), while the numbers we determined for several key fission yeast proteins were 1.5-2 times lower than previously reported (Figure 2E, 2F and 2G) (Sirotkin et al., 2010). In total, our direct, side-by-side comparison indicates that the numbers of proteins at endocytic sites in these two yeasts are much more similar than was previously thought. The modest differences between the two yeasts may help to explain the differences in the dynamics reported in the following sections.

### Particle tracking of fluorescent protein-tagged endocytic proteins reveals different behaviors during internalization in budding and fission yeast

In contrast to the budding yeast, the morphological details of fission yeast endocytic vesicle formation are not well defined. To elucidate such information for fission yeast, we next compared the recruitment and spatial dynamics of endocytic proteins side by side in budding and fission yeast. We acquired images in the equatorial plane of cells at high frame rate (more than 7Hz). The time-lapse images were then processed and analyzed using the ImageJ Particle Tracker plugin, which records the centroid position and fluorescence intensity of a fluorescently-tagged protein over time to generate a trajectory (Figure 1-figure supplement 3). We developed Python-based software to further analyze the trajectory data.

The dynamics of the actin filament network were compared in the two yeast species as a function of time by tracking scAbp1-GFP (Actin-Binding Protein 1) in budding yeast and spFim1-GFP (fimbrin) in fission yeast in data collected from the same movies at a frame rate of 9 Hz (Figure 3A). Both scAbp1-GFP and spFim1-GFP patches show little motility at the beginning of their lifetimes in endocytic patches while the fluorescence intensity gradually increases (Figure 3B and 3C). All trajectories for each protein were aligned at 50% of their maximum intensity, then averaged and the average plotted (Figure 3D and 3E). The scAbp1-GFP moves about 125 nm with relatively high regularity (as indicated by the small standard deviation, Figure 3-figure supplement 1A) while the intensity approaches the maximum (Figure 3D). Once the scAbp1-GFP intensity starts to decline, its movement becomes far more irregular (as indicated by a pronounced increase in the standard deviation) (Figure 3D and Figure 3-figure supplement 1A). Previous EM studies suggested that in budding yeast the distance between the base and the tip of the invaginated endocytic membrane often reaches about ∼100 nm in depth before vesicle scission occurs (Kukulski et al., 2012). Considering the fluorescence and EM data that reflects on endocytic membrane morphology, we conclude that the transition to irregular movement of scAbp1-GFP beyond ∼125 nm from the origin likely represents endocytic vesicle release in budding yeast. Our data for budding yeast agree well with those of Picco et al. (Picco et al., 2015), in which Rvs167 (budding yeast amphiphysin) dynamics was used to predict the scission point. Thus, the quantitative analysis using our custom software appears to indicate the timing of the full endocytic progression from site formation to invagination initiation to vesicle release.

**Figure 3.**
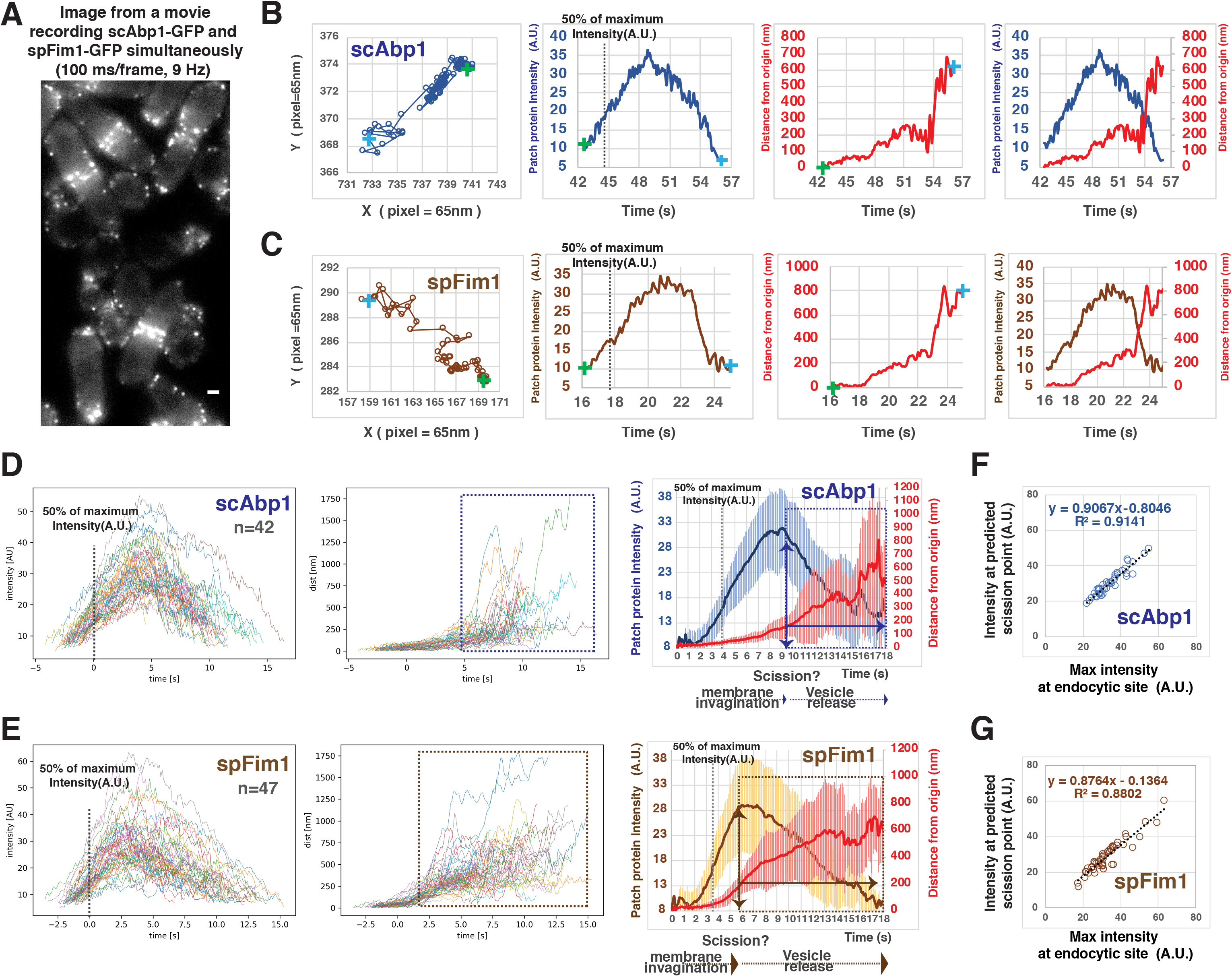
Quantitative comparison of endocytic actin dynamics in budding and fission yeast. A, Single frame from a movie of simultaneously imaged *scABP1-GFP* in budding yeast and *spFIM1-mGFP* in fission yeast. B and C, Single endocytic events represented by scAbp1 in budding yeast (B) and by spFim1 in fission yeast (C) were tracked and then analyzed using our custom software. Graphs from left to right: (Left) Protein patch centroid position over time. Consecutive positions are connected by lines. “+” in green or in blue indicates the first or the last position, respectively. (Left Center) Fluorescence intensity over time. Dotted line indicates the time point when fluorescence intensity reaches 50% of its maximum intensity. (Right Center) Displacement over time. Displacement from the origin is the distance from the position of each time point to the starting position (“+” in green). (Right) Fluorescence intensity and displacement over time. D and F, Numerous endocytic events tracked by imaging fluorescent scAbp1 (D) or spFim1 (E) were analyzed and aligned. Graphs from left to right: numerous endocytic events aligned to the point of 50% of maximum intensity (indicated by dotted line); displacement data for same cells as in right panel aligned according to 50% maximum intensity point (boxed areas represent inferred movement after scission); combined average results from graphs on the left (note that time is rescaled on the averaged data graphs). Dotted line indicates the time point when fluorescence intensity reaches to 50%. Vertical line with arrow indicates the inferred moment of scission predicted by the dramatic increase in standard deviation (Figure 3-figure supplement 1). Standard deviation is represented by the shadow around the average line. F and G, Inferred endocytic scission is tightly correlated with the time when endocytic actin assembly reaches its maximum for budding (F) and fission yeast (G). The scale bars on cell pictures are 2µm.

The dynamics of spFim1-GFP (Figure 3E) followed a similar course as that for scAbp1-GFP (Figure 3D). Strikingly, however, the irregular movement of spFim1-GFP (indicated by a sharp increase in the standard deviation, Figure 3-figure supplement 1B) occurs after its initial regular displacement reaches ∼200 nm. Since the irregular movement likely reflects movement of the free vesicle, endocytic membrane invaginations in fission yeast appear to be substantially deeper than in budding yeast before vesicle scission occurs. Moreover, in both yeasts, the predicted scission time point is tightly correlated with the maximum intensity of the endocytic actin network (Figure 3F and 3G).

To independently validate the conclusions reached from the actin network dynamics observations, we carried out a similar quantitative analysis for the endocytic coat proteins scSla1-GFP and spPan1-GFP in budding and fission yeast, respectively (Figure 4). In both yeasts the coat proteins stay non-motile for a relatively long period (∼30 s –120 s) and then begin to move (Figure 4A and 4B). We defined the initiation of coat protein movement by an inflection point in the “distance from origin” vs “time” graph (Figure 4A and 4B). Our custom software automatically detects the inflection point for each trajectory and aligns multiple trajectories at the inflection point (Figure 4C and 4D). After the inflection point, scSla1-GFP and spPan1-GFP move regularly (indicated by a relatively small standard deviation) for about 4-5 s, followed by a sharp transition to a period of irregular movement (indicated by a pronounced increase in the standard deviation) (Figure 4C and 4D). Thus, we conclude that endocytic vesicle scission likely occurs around 4-5 s after the initiation of endocytic patch movement in both yeasts. The scSla1-GFP patches in budding yeast move ∼100nm before vesicle scission (Figure 4C). Similar results were obtained for scSla2-GFP, scPan1-GFP and scEnd3-GFP (data not shown). However, in fission yeast, spPan1-GFP (Figure 4D) and spSla2-GFP (data not shown) move ∼200 nm during the 4-5 s after the initiation of coat protein movement and before the transition to the irregular movement phase, further supporting the conclusion that the fission yeast endocytic membrane invagination reaches a greater depth prior to vesicle scission than in budding yeast. While the initiation of both spPan1-GFP and scSla1-GFP patch movement is tightly correlated with attainment of the maximum intensity for each protein (Figure 4E), the invagination speed in fission yeast is about twice as fast as in budding yeast (51.8 nm/s vs 23.8 nm/s) (Figure 4F).

**Figure 4.**
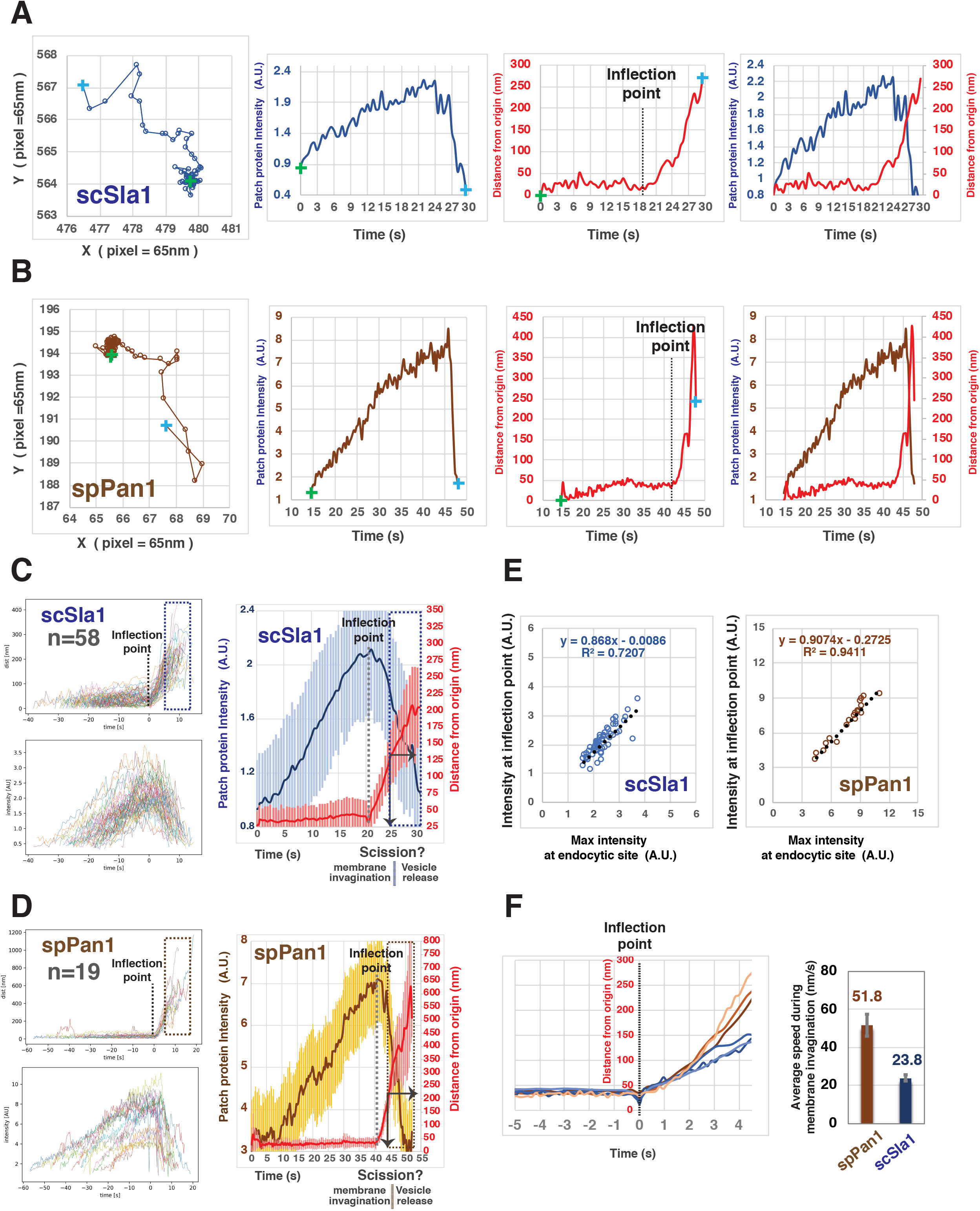
Quantitative comparison of endocytic coat dynamics for budding and fission yeast. A and B, Single endocytic event detected by tracking fluorescent scSla1 in budding yeast (A) or spPan1 in fission yeast (B). Graphs from left to right: protein patch centroid position over time, “+” in green or in blue indicate the first and last positions, respectively; fluorescence intensity over time; displacement over time (“+” in green and blue indicate starting and end points, respectively while the dotted line indicates the inflection point at which time the coat protein begins to move; fluorescence intensity and displacement over time. C and D, Numerous endocytic events detected by tracking fluorescent scSla1 (C) or by spPan1 (D) were analyzed and aligned. Graphs from left to right: numerous endocytic events aligned at inflection points indicated by the dotted line, the boxed area represents movement after inferred scission event; fluorescence intensity aligned by the movement inflection point; combined average results from the two graphs on the left (note that time is rescaled in the average graph and that the dotted line indicates inflection point), vertical line with arrow indicates the moment of scission predicted by the dramatic standard deviation increase. Standard deviation is represented by the shadow around the average line. E, Initiation of endocytic membrane invagination is tightly correlated with the time when the endocytic coat reaches its maximum amount in budding and fission yeast. F, Speed of coat protein movement during the inferred invagination process prior to scission. Each of the brownish or bluish lines represents the average displacement of spPan1 or scSla1, respectively, over time from one experiment. Three independent experiments were performed for the indicated proteins. Bar graphs show rates for coat proteins.

Additionally, we examined patch dynamics for both spPan1-GFP and spFim1-mCherry at a rate of more than 7 frames per second in fission yeast (Figure 5A and 5B, Figure 5-figure supplement 1). Actin assembly (imaged using spFim1-mCherry) begins when spPan1-GFP reaches its peak intensity (Figure 5B, dotted line 1). About ∼2 s later (Figure 5B, dotted line 2), both fluorescent proteins start to move in a regular manner more than 200 nm over 4-5 s prior to beginning to move irregularly when the endocytic vesicle is presumed to be released. At this point the spFim1-mCherry intensity peaks (Figure 5B, dotted line3 and Figure 5-figure supplement 1).

**Figure 5.**
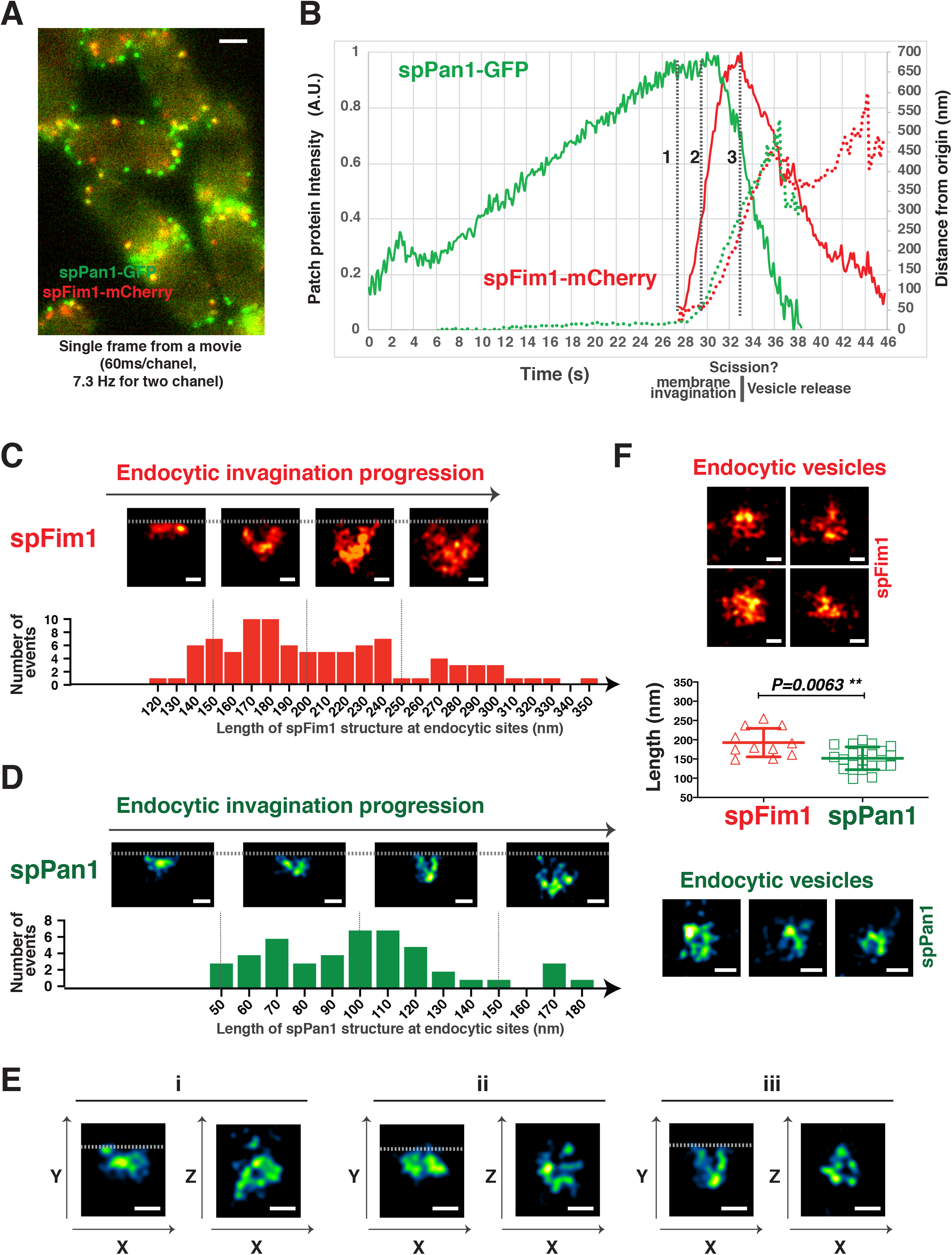
Spatial-temporal relationship and 3D-STORM imaging of coat proteins and the actin network at fission yeast endocytic sites. A, Single frame from a movie of *spPan1-GFP spFIM1-mCherry* expressed in fission yeast cells. B, Alignment of average intensity (solid lines) and displacement (dotted lines) of spPan1-GFP (coat) and spFim1-mCherry (actin cytoskeleton) patches (n=15) (Figure 5-figure supplement 1 for details). Vertical dotted line 1 indicates the time when actin assembly is first detected. Dotted line 2 marks initiation of membrane invagination. Dotted line 3 indicates inferred scission event. C and D, The length of nanoscale structures of spFim1 (n=92) or spPan1 (n=46) at endocytic sites revealed by STORM analysis was measured and plotted. Representative STORM images of different length and quantitative analysis of spFim1 (C) and spPan1 (D). The histogram shows the frequency distribution of observed endocytic structure lengths. E, 3D-STORM image analysis reveals ring-like Pan1 organization in the XZ dimensions. (F), Representative images of presumed endocytic vesicles in the cytoplasm. The length of longest axis of each presumed vesicle was measured and plotted. Fim1-labled structures were pseudo-colored red hot and Pan1-labled structures were pseudo-colored green fire blue (C-F). Scale bars on cell pictures (A) are 2µm. The scale bars on STORM images (C, D, E and F) are 100nm.

Together, our side by side quantitative analysis revealed intriguing unexpected differences in endocytic invagination development between budding and fission yeast. The results demonstrate that both yeast spend similar time deforming the membrane prior to scission. However, the resulting endocytic invagination of fission yeast appears to be nearly twice as long as in budding yeast.

### Three-dimensional stochastic optical reconstruction microscopy (3D-STORM) reveals novel nanoscale structural details of endocytic vesicle formation in fission yeast

To acquire a nanoscale structural description of endocytic site formation in fixed fission yeast cells, we carried out 3D-STORM analysis by labeling GFP-tagged endocytic proteins with AF647-conjugated anti-GFP nanobodies in fixed cells (Kaplan and Ewers, 2015; Mund et al., 2014). Nanobodies conjugated to the AF647 are small enough to readily penetrate the yeast cell wall and thus eliminate the requirement for its prior enzymatic removal, allowing high quality structural preservation and reduced localization error.

Numerous endocytic sites labeled by spFim1-GFP or spPan1-GFP in fixed cells were examined at the equatorial plane of the cells by 3D-STORM (Figure 5C-5F, Figure 5-figure supplement 2), so invaginations of the plasma membrane were in the image plane. spFim1 was used to label the endocytic actin network. The length of the spFim1 structures ranged from 120 nm to 350 nm (Figure 5C), which likely represents progression through the endocytic pathway. The size of these structures is consistent with the spatial distribution of single molecule localizations of Fim1p-mEOS3.2 observed for fission yeast endocytic actin clouds by live-cell super resolution microscopy (Arasada et al., 2018). In a similar STORM study in fixed budding yeast, the length of the endocytic actin network was estimated range from 70 nm-240 nm (Mund et al., 2018). EM studies in budding yeast suggest that a dense network of actin filaments occupies a large area outside the invaginated membrane prior to scission. This area is expected to be larger than that traveled by the centroid of actin network fluorescence in the dynamics experiments described above because those studies do not describe the extent of the network but instead its center. Thus, the greater length of endocytic actin structures observed in fission yeast is consistent with the possibility that the invaginated membrane that it covers is deeper than in budding yeast. The length of spPan1 structures in fission yeast ranged from 50 nm to 180 nm (Figure 5D), which is much smaller than the range for spFim1 structures. spPan1 is a coat protein that is expected to localize on the invaginated membrane, more precisely representing the morphology of invaginated membrane. In addition, 3D-STORM analysis revealed that spPan1 forms a ring structure in the X-Y plane, presumably encircling the invaginated membrane (Figure 5E). A similar ring structure could also be observed with the spFim1-GFP labeling (Figure 5-figure supplement 2D). Moreover, in the equatorial plane we observed structures that we interpret as pinched-off endocytic vesicles based on their distances of >500 nm from the cell cortex (Figure 5F). The average long axis of endocytic vesicles labeled by spFim1 and by spPan1 was around 193±37 nm or 152± 30 nm, respectively. A similar nanoscale structural analysis is currently not available for budding yeast endocytic vesicles.

Together, our 3D-STORM results further support the conclusion that fission yeast endocytic invaginations are deeper than those observed in budding yeast.

### Spatial-temporal relationship between two major Nucleation Promoting Factors (NPFs) and the actin network during fission yeast endocytosis

Our results thus far identified unexpected novel features of fission yeast endocytic invagination that differ from those in budding yeast. Thus, it is necessary to reevaluate the mechanism of CME actin force generation that promotes membrane invagination in fission yeast. We next performed a comparative study of the actin assembly mechanism that drives endocytic internalization in the two yeasts. Our studies focused on WASp (spWsp1) and type I myosin (spMyo1), which in fission yeast are the two major NPFs that activate the Arp2/3 complex at endocytic sites (Sirotkin et al., 2005).

scLas17 in budding yeast has been reported to remain near the base of the pit (close to the plasma membrane) during endocytosis while the actin network moves into the cytoplasm (Kaksonen et al., 2003). In contrast, spWsp1 in fission yeast was reported to move away from the base, similar to the actin network (Sirotkin et al., 2010). One possible source of the different observations on these two yeasts is that a fluorescent protein-tag was added at scLas17’s C-terminus in budding yeast but was added at spWsp1’s N-terminus in fission yeast. We thus generated a budding yeast strain in which the GFP tag was added at the N-terminus of Las17 (Figure 6-figure supplement 1) by making an in-frame fusion at the native *LAS17* locus. More than 70% of GFP-Las17 patches showed similar dynamics as observed for Las17-GFP (Figure 6-figure supplement 1A and 1B). However, some GFP-Las17 patches appeared to split into two parts toward the end of the patch lifetime: the majority of the GFP-Las17 signal remained non-motile until it disappeared from the endocytic site while a small amount of the GFP-Las17 moved away from the cell surface (Figure 6-figure supplement 1A). In spite of these differences, the populations of actin patches in cells depending on GFP-Las17 and Las17-GFP were indistinguishable by two criteria: (1) patch lifetimes were identical (Figure 6-figure supplement 1B); (2) the actin network markers Abp1p-RFP and Myo5p-RFP had similar spatial-dynamic behaviors in a *GFP-LAS17* strain (Figure 6-figure supplement 1C) and a *LAS17-GFP* strain (Sun et al., 2006).

**Figure 6.**
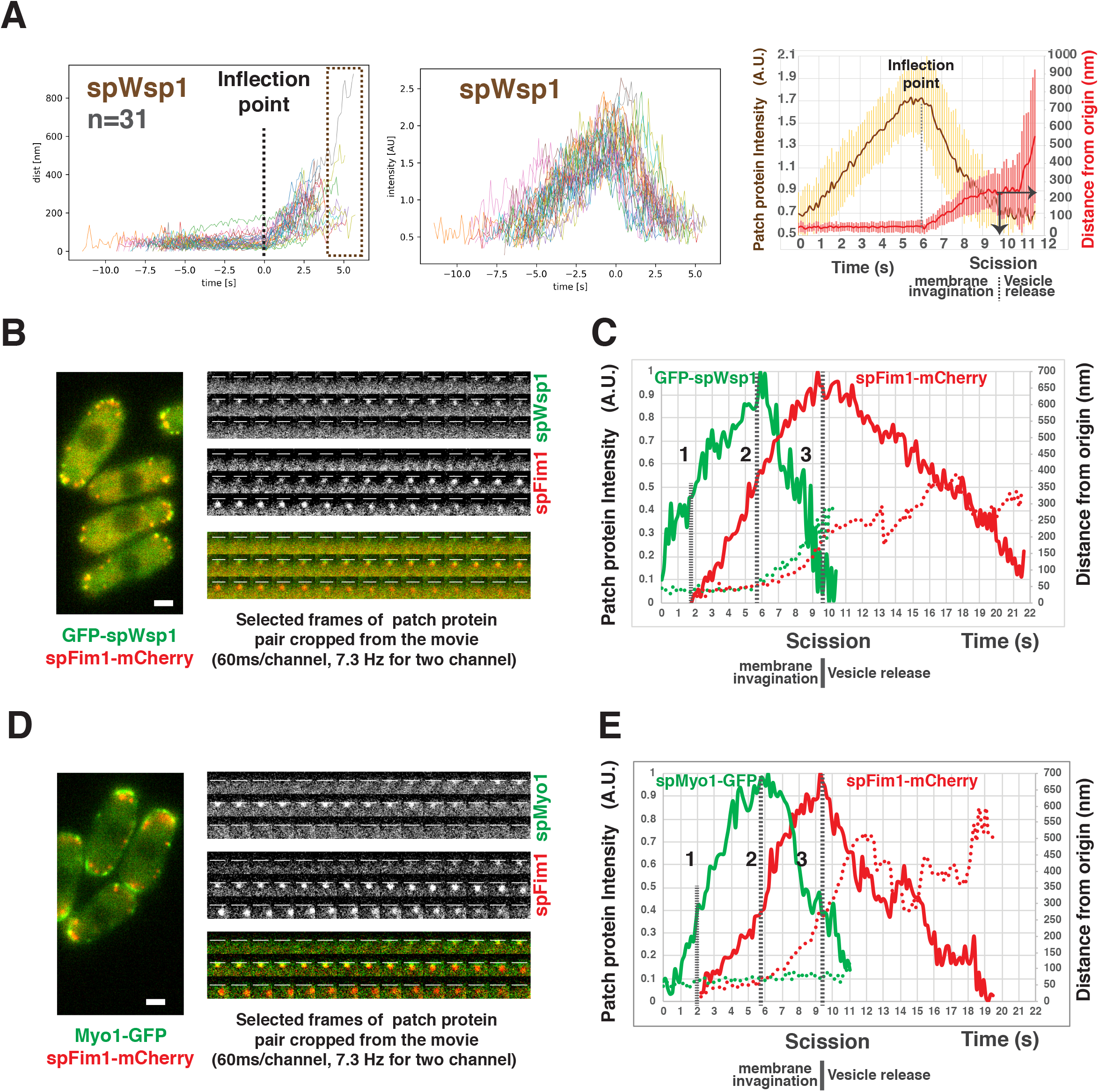
Spatio-temporal relationship between two major NPFs and endocytic actin assembly in fission yeast. A, Dynamics of spWsp1. Numerous endocytic events represented by GFP-spWsp1 were analyzed and aligned. Graphs from left to right: (right) Protein position vs time for endocytic events (n=31) aligned at inflection points indicated by the dotted line. The boxed area represents movement after presumed scission event. (middle) Fluorescence intensity aligned on basis of alignment in right graph. (left) Averaged results from data in two graphs on the left. Note that time is rescaled in this graph. The dotted line indicates inflection point. Vertical solid line with arrow indicates the moment of scission predicted by the dramatic standard deviation increase. B, Single frame from a movie of fission yeast expressing *GFP-spWSP1 spFIM1-mCherry* (left panel). Time series showing composition of a single endocytic sites (right panels). C, Alignment of average intensity and displacement for GFP-spWsp1 and spFim1-mCherry patches (n=8) (Figure 6-figure supplement 2 for details). Dotted line1 indicates actin assembly initiation. Dotted line 2 indicates initiation of membrane invagination. Dotted line 3 indicates inferred scission. D, Single frame from a movie of fission yeast expressing *MYO1-GFP spFIM1-mCherry* (left panel). Time series showing composition of a single endocytic sites (right panels). E, Alignment of average intensity and displacement for GFP-spWsp1 and spFim1-mCherry patches (n=9) (Figure 6-figure supplement 3 for details). Dotted line 1 indicates actin assembly initiation. Dotted line 2 indicates initiation of membrane invagination. Dotted line 3 indicates inferred scission. Scale bars are 2µm.

On the other hand, our high frame rate live-cell image analysis confirmed that GFP-spWsp1 patches move away from their origin, into the cytoplasm in fission yeast (Figure 6A). Intriguingly, our newly developed alignment software revealed that GFP-spWsp1 only begins to move when it accumulates to its maximum level (Figure 6A). GFP-spWsp1 patches then move ∼200nm over ∼4s, while their fluorescence intensity rapidly declines to the baseline level (Figure 6A). The regular phase of GFP-spWsp1 movement prior to the onset of the irregular movement corresponds well with the endocytic invagination dynamics observed with other fission yeast proteins (Figure 3, Figure 4, and Figure 5). Our results (Figure 6-figure supplement 1 and Figure 6A) also concluded that the differences between WASp dynamics in the two yeasts are not the result of differences in the fluorescent protein fusions employed or differences in the analytical methods used in different studies.

Previous studies using single color imaging at low frame rates (0.3-0.5Hz) suggested that the two major fission yeast NPFs (spWsp1 and spMyo1) appear at endocytic sites a few seconds before (Arasada and Pollard, 2011), or with similar timing to (Sirotkin et al., 2010), onset of actin assembly. To better understand how the two NPFs regulate endocytic actin assembly, we investigated the spatiotemporal relationship between the Fim1 and spWsp1 or spMyo1 using high time resolution (7.3 Hz) two-color imaging (Figure 6B-6E, Figure 6-figure supplement 2, Figure 6-figure supplement 3). GFP-spWsp1 appears at the cell cortex first, then actin assembles (spFim1-mCherry) only when GFP-spWsp1 accumulates to approximately 50% of the maximum level (Figure 6C, dotted line 1). Once GFP-spWsp1 reaches the maximum level, both GFP-spWsp1 and the actin network marker spFim1 begin to move (Figure 6C, dotted line 2). While GFP-spWsp1 and the actin network move ∼200 nm from the origin, the GFP-spWsp1 signal declines to the baseline level while the actin network approaches the maximum level (Figure 6C, dotted line 3). Consistently, vesicle scission appears to occur when actin assembly reaches its maximum level (Figure 3G, Figure 5B, Figure 6C, and Figure 6-figure supplement 2). spMyo1-GFP accumulates at endocytic sites with similar timing to GFP-spWsp1, using spFim1-mCherry accumulation as a temporal alignment reference (Figure 6D, 6E and Figure 6-figure supplement 3). However, as in budding yeast (Sun et al, 2006), spMyo1-GFP stays non-motile during its lifetime while the actin network moves into the cytoplasm (Figure 6D and 6E).

We conclude that actin assembly is initiated when both spWsp1 and spMyo1 reach approximately 50% of their peak levels in fission yeast. Importantly, spWsp1 and spMyo1 are recruited to their maximum levels before the membrane detectably begins to invaginate, supporting the conclusion that spWsp1 and spMyo1 both promote actin nucleation to initiate membrane deformation. Moreover, the rapid disassembly of spWsp1 and spMyo1 from endocytic sites supports the notion that actin nucleation by the two NPFs rapidly decreases while the invagination elongates.

### The relative importance of spWsp1 and spMyo1 NPF activity for fission yeast endocytosis

The previously proposed two-zone model for fission yeast implies that spWsp1 and spMyo1 define two independent pathways for actin assembly at the endocytic sites to facilitate endocytic invagination (Arasada and Pollard, 2011) (Figure 1-figure supplement 1). Deletion of either Myo1 or Wsp1 stops endocytosis (Arasada and Pollard, 2011) but removes not only their NPF activity but also other functions (Pedersen and Drubin, 2019; Sun et al., 2006) provided by domains that do not interact with the Arp2/3 complex. To specifically test the importance of the NPF activity, we created mutants of Myo1 and Wsp1 only lacking the carboxy terminal “CA” motifs that bind and activate Arp2/3 complex.

The spwsp1CAΔ-GFP mutant protein accumulates and persists at endocytic sites nearly 3 times longer than the wild-type protein without detectable movement (Figure 6A, Figure 7A, and 7B). Two-color live cell imaging demonstrated that the actin network (spFim1-mcherry) still assembles, but that most of the assembled actin is non-motile during its life-time, indicating that endocytic internalization is largely impaired in the *spwsp1-CAΔ* mutant (Figure 7C and 7D, Video1, and Video 2). In contrast, endocytic internalization occurs rather regularly in the *spmyo1CAΔ* mutant (Figure 7E and 7F, Video3, and Video 4), in which its “CA” motif was precisely truncated at its genomic locus. The myo1CAΔ-GFP protein exhibits comparable lifetimes to wild-type spMyo1-GFP (Figure 7B).

**Figure 7.**
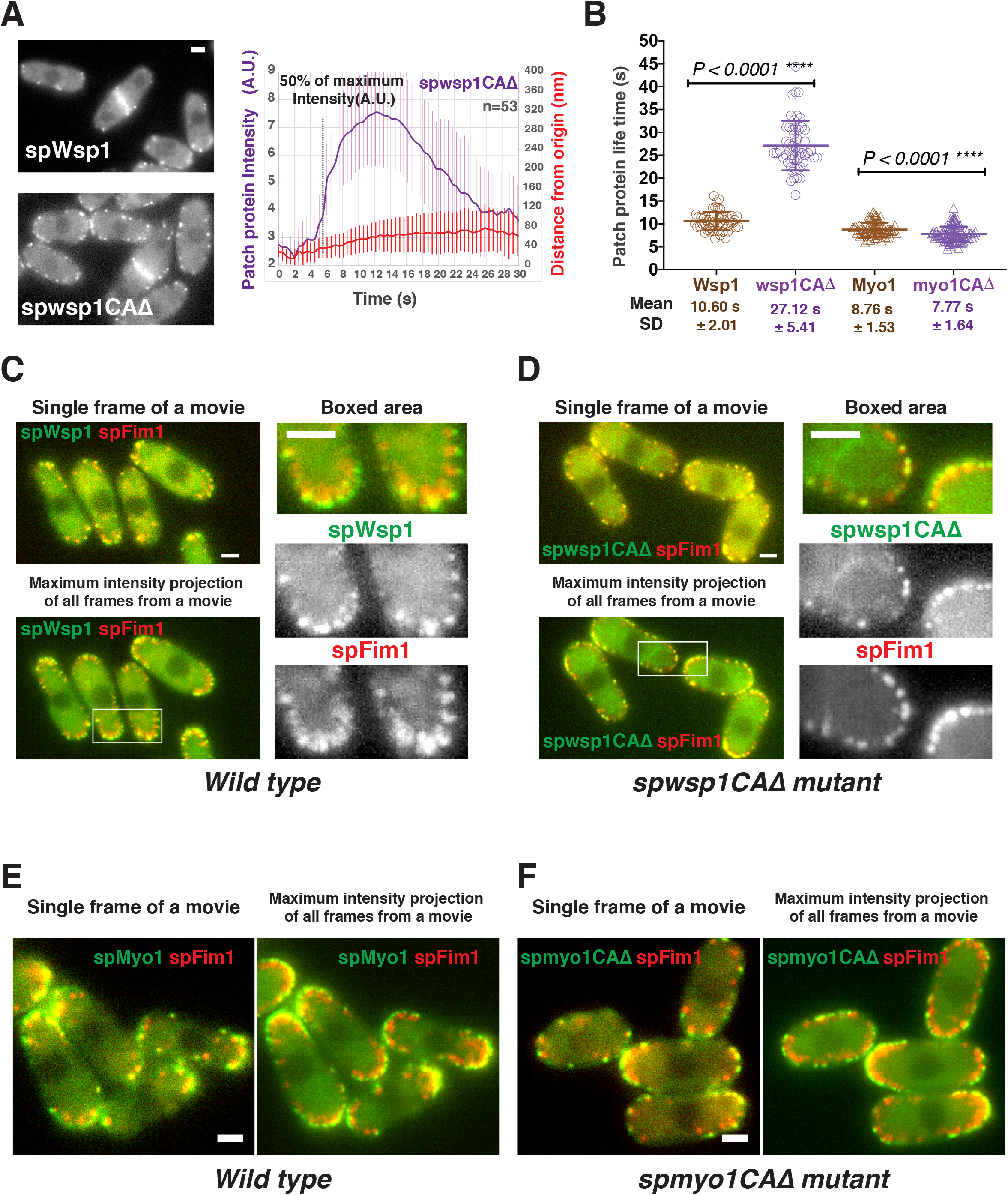
Functional analysis of CA domains in two fission yeast nucleation promoting factors. A, Dynamics of spwsp1CAΔ-GFP patches. Single frame from movie of cells expressing *GFP-spWSP1* (left upper panel) or *spWSP1-CAΔ-GFP* cells (left bottom panel). Alignment of average intensity and displacement for spwsp1-CAΔ-GFP patches (n=53) (right panel). Dotted line indicates 50% maximum fluorescence intensity. B, Patch protein life times for indicated proteins. C and D, Endocytic internalization is severely impaired in *spwsp1-CAΔ* mutants. A single frame (upper left) and Maximum intensity projection of all frames (lower left) from a movie of cells expressing *GFP-spWSP1 spFIM1-mCherry* (C) or *spwsp1-CAΔ-GFP spFIM1-mCherry* (D). The maximum intensity projections reveal extent of patch protein movement over time. Enlarged views of the boxed-areas shown in left panels are shown on right. E and F, Single frame (left) and maximum intensity projection (right) from of all frames of movies for *MYO1-GFP spFIM1-mCherry* cells (E) or *spmyo1-CAΔ-GFP spFIM1-mCherry* cells (F). Scale bars are 2µm.

Together, these observations indicate that the spWsp1 NPF activity is essential for endocytosis, that the spMyo1 NPF activity is not, and they suggest that two-zone actin assembly is not required for endocytosis in fission yeast.

## Discussion

This study quantitatively compared several important aspects of CME in distantly related yeasts to gain evolutionary insights into underlying mechanisms and how they can be adapted for the specific demands of different organisms and cell types in multicellular organisms.

### Quantitation of endocytic protein numbers

Knowing how many of each type of protein is present at an endocytic site is crucial for developing a quantitative understanding of CME. Previous studies reported larger numbers (up to 5 fold) of endocytic proteins in fission yeast relative to homologues in budding yeast (Picco et al., 2015; Sirotkin et al., 2010) (Figure 1-table supplement 1). However, these counting studies were done in several different laboratories, which focused exclusively on either budding or fission yeast, and they largely used different experimental methods. In fact, different results have been reported for the number of molecules of the same protein in the same yeast species (Figure 1-table supplement 1) (Basu and Chang, 2011; Chen and Pollard, 2013; Epstein et al., 2018; Galletta et al., 2012; Picco et al., 2015; Sirotkin et al., 2010). To reconcile these differences, we made great efforts to validate our methods for an accurate quantitative analysis of protein numbers, and to make comparisons between the yeasts directly.

Firstly, to determine the relative abundance of homologous proteins in the two yeasts, we imaged mixtures of budding and fission yeast cells on the same microscope field. Thus, most potential sources of inconsistencies were eliminated. The imaging condition we established should also be useful for comparative studies of other cellular processes between the two yeast species. The fluorescence intensities of homologous proteins in the two yeasts were determined in the same movie, which was processed and analyzed by exactly same procedure, further ensuring a faithful comparison. For these reasons, we believe that our approaches represent an improvement in accuracy over those used previously. Strikingly, we determined that endocytic protein homologues in the two yeasts are more similar in abundance than indicated by numbers available in the published literature (Figure 1, Figure 2A, Figure 1-table supplement 1).

Secondly, we adopted previously developed 120 sfGFP nanocages (Hsia et al., 2016) as a new fluorescent standard for counting absolute protein numbers. Each nanocage is a stable icosahedron with a diameter of ∼25 nm. The nanocages appear as diffraction limited spots comparable to endocytic patch proteins resolved by 2D-wide-field fluorescence microscopy. The nanocages are easy to purify and handle and as a result, large sample sizes can be imaged in the same field, giving more reliable and precise results providing that the medium has a pH of ∼7.0 (details in Experimental Procedures). Moreover, intensity histograms for 120-sfGFP-tagged nanocages prepared on different days give highly reproducible results (Figure 2-figure supplement 1). As validation for our new standard, we used the 120-sfGFP tagged nanocages to estimate that 16-kinetochore clusters carry 92±19 Cse4-sfGFP molecules, consistent with results of several previous studies (Coffman et al., 2011; Galletta et al., 2012; Lawrimore et al., 2011). Thus, we established that extracellular nanocages with 120 sfGFPs serve as an easy-to-use and reliable standard for accurate determination of absolute numbers of proteins.

Our counts of Abp1 and Arp2/3 complex in actin patches of budding yeast agree well with those of Galletta et al. (Galletta et al., 2012) (Figure 1-table supplement 1), who used centromere protein Cse4 as an internal standard. We also agree with Picco et al. (Picco et al., 2015) on the relative abundances of seven budding yeast endocytic proteins but our counts are mostly ∼2 fold higher than theirs (Figure 2-figure supplement 2, Figure 1-table supplement 1). These systematic differences in absolute number estimates are likely due to use of different calibration standards. Picco et al. used the kinetochore protein Nuf2 at the anaphase to telophase transition as a standard. However, a recent study demonstrated that Nuf2 levels change dynamically during the anaphase to telophase transition (Dhatchinamoorthy et al., 2017), which complicates its use a standard. For fission yeast, our counts for some of the endocytic proteins (Fim1, spWsp1 and spMyo1) are substantially lower than some (Epstein et al., 2018; Sirotkin et al., 2010) but not all (Arasada and Pollard, 2011; Chen and Pollard, 2013) previous measurements (Figure 1-table supplement 1). These differences might be explained by the live cell imaging conditions and methods to correct for local background (Figure 1-figure supplement 3).

To date, molecular numbers of endocytic mammalian proteins at CME sites have mostly not been determined due to technical difficulties associated with genetic manipulation of genomes in animal cells. However, dynamin numbers have been determined in genome-edited mammalian cells (Cocucci et al., 2014; Grassart et al., 2014). Pertinent to the current study, Akamatsu et al (submitted) recently generated a cell line that expresses an endogenously-tagged fluorescent Arp2/3 subunit and estimated its peak number at endocytic sites as ∼200, which is lower than, but close to, the number found at budding (294) and fission (302) yeast CME sites (Figure 2G). Thus, the numbers of endocytic proteins at CME sites in yeast may predict numbers in human cells, and differences may provide a basis for understanding cell type-specific requirements for CME such as the forces necessary for vesicle formation.

### Comparison of protein dynamics and endocytic site morphology for CME internalization stages

In this study, we analyzed fission yeast endocytic internalization using high time resolution live-cell imaging and custom analytical software. Wide-field fluorescence microscopy in the equatorial plane of yeast cells enabled fast acquisition, high signal intensity and the opportunity to observe the complete endocytic internalization process from a side view. However, the endocytic protein imaged by this method may not remain in the same focal plane during the full-time course, especially after endocytic vesicle scission. For this reason and because of variation in kinetic profiles of different events, the fluorescence intensity profiles varied from one recorded trajectory to another. We therefore developed a conceptually different alignment method from those used previously (Berro and Pollard, 2014; Picco et al., 2015) to suit our imaging method. Our custom software automatically transformed the tracking data for numerous individual events into visually intuitive graphs, showing the 2-D trajectory, raw (unprocessed) fluorescence intensity, and displacement over time. Comparing graphs for numerous events allowed us to determine the signatures for each endocytic protein and align the trajectories of the proteins. Our software not only recapitulates the dynamic endocytic protein profiles reported previously (Berro and Pollard, 2014; Picco et al., 2015), but also distinguishes the stages of endocytic internalization, such as invagination elongation, scission, and vesicle release.

In the early studies of fission yeast, the trajectories of most endocytic protein(s) were aligned relative to the initiation of patch movement in movies of lower time resolution (0.3-0.5 Hz) (Arasada and Pollard, 2011; Sirotkin et al., 2010) without knowing how this movement correlates with specific endocytic steps. A later study improved the temporal resolution of Fim1 recruitment (Berro and Pollard, 2014; Picco et al., 2015), but knowledge of how the resulting timeline maps onto morphological stages of CME was still unclear. Our high time resolution movies (more than 7 Hz) and alignment software revealed that the centroid position of both the endocytic actin network (represented by spFim1) and the endocytic coat (represented by spPan1) move ∼200 nm from their initiation sites prior to predicted scission (Figure 3, Figure 4, and Figure 5A and 5B). These results suggest that the endocytic membrane invagination is ∼200 nm deep at the time of scission, nearly twice the depth observed for budding yeast. The nanoscale structural organization of spFim1 and spPan1 at endocytic sites determined by super-resolution imaging further supports this interpretation (Figure 5C, 5D). Moreover, the initiation of membrane invagination is tightly correlated with attainment of the maximum intensity for spPan1-GFP, while the predicted scission time point is tightly correlated with the maximum intensity of Fim1 (Figure 3E and 3G, Figure 4D and 4E, Figure 5 A and 5B). Our results agree with and provide an explanation for the previous finding that the motion of Fim1 patches is diffusive rather than directional when the peak number at endocytic sites is reached (Berro and Pollard, 2014). Another intriguing result is that the elongation speed for membrane invagination is twice as fast in fission yeast as in budding yeast (Figure 4F).

The two yeasts studied here show dramatic differences in endocytic site dimensions and dynamics. As similar differences have been reported within a range of mammalian cell types (Dambournet et al., 2018; McMahon and Boucrot, 2011), comparative studies between the yeasts might identify mechanisms for controlling CME structure and dynamics more generally.

### “Push and pull” vs “two-zone” models for force production by actin polymerization

A two-zone model, in which distinct zones of actin filament nucleation mediated by type I myosin and WASp generate two separate actin networks that push against each other, was proposed to generate endocytic forces in fission yeast (Arasada and Pollard, 2011) (Figure 1-figure supplement1). However, when we eliminated the actin filament nucleation promoting activity of spMyo1, spWsp1 alone was sufficient to support robust endocytic internalization with only one zone of actin assembly (Figure 7F). Thus, the two-zone mechanism (Figure 1-figure supplement1) is apparently not essential for fission yeast endocytosis. In addition, we found that membrane deformation starts when the numbers of spWsp1 and spMyo1 peak at endocytic sites (Figure 6A) supporting the conclusion that spWsp and spMyo1 promote actin nucleation to initiate membrane deformation. Interestingly, both yeasts accumulate similar maximum numbers of Arp2/3 complexes (∼300) and HIP1R homologs (scSla2 and spEnd4, ∼130) at endocytic sites (Figure 2E and 2F), suggesting similar capacities for producing actin filament branches and for connecting the actin network to the endocytic coat. Thus, at least to some extent, the push and pull mechanism proposed for budding yeast (Figure 1-figure supplement1) is sufficient for fission yeast CME, consistent with a recent models of force production by actin assembly (Nickaeen et al., 2019).

While in mammalian cells the detailed dynamics of N-WASp and type I myosin during CME membrane deformation are not clear, both localize at mammalian CME sites (Krendel et al., 2007; Merrifield et al., 2004). However, mammal type I myosin does not carry CA motif, and so it is not likely to directly activate the Arp2/3 complex. Evidence presented here indicates that two-zones of actin assembly may not be required for mammalian CME.

### Conclusions from quantitative, evolutionary comparison of CME in distantly related yeasts

Integrating our quantitative comparative analysis of endocytosis with previous knowledge revealed several key similarities and differences in budding yeast and fission yeast in terms of molecular numbers, architecture and function of the endocytic coat and associated actin machinery (Figure 8).

**Figure 8.**
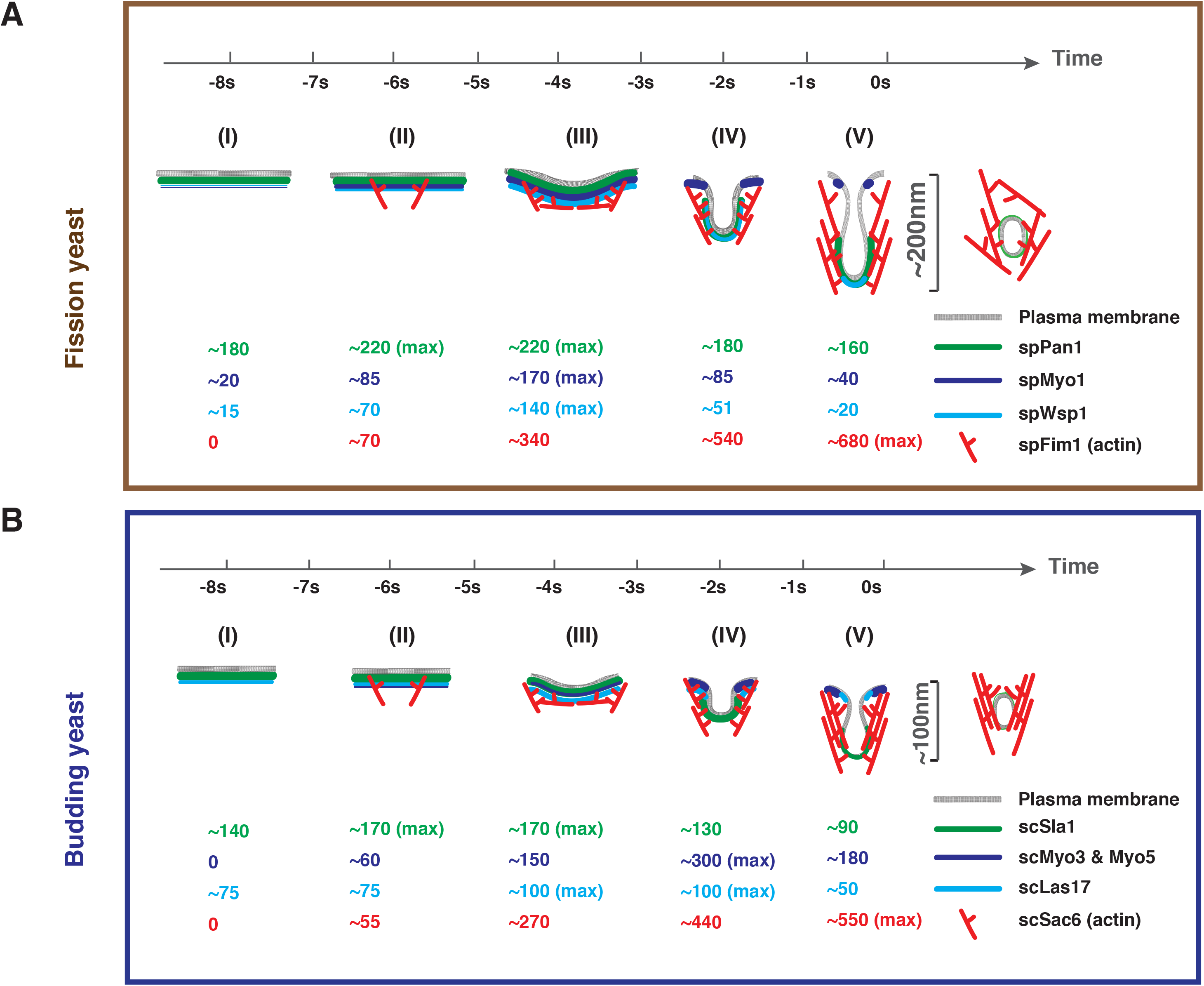
Comparison of endocytic vesicle formation in fission and budding yeast. Timeline and summary of the average molecule numbers for indicated coat proteins and actin machinery components in fission (A) and budding yeast (B) endocytosis. Scission occurs at the time 0. Steps (I) through (V) represent the full process from membrane invagination initiation to vesicle scission. See text for detailed discussion.

The first conclusion is that the two yeasts trigger actin assembly at endocytic sites similarly. The homologous proteins spPan1 and scSla1 reach their maximum numbers at endocytic sites before actin assembly is initiated (Figure 4C, 4D, 4E and Figure 8A(II), 8B(II)). Since actin assembly is required to generate membrane curvature in yeast (Kukulski et al., 2012), this observation supports the hypothesis that the endocytic coat first assembles as a planar structure that subsequently deforms into a tubular invagination, as also proposed for CME in mammalian cells (Avinoam et al., 2015). Recruitment of spWsp1 at endocytic sites in fission yeast begins as spPan1 peaks (Figure 5B and Figure 6C, Figure 8A(I)) followed in both yeasts by switch-like activation of the Arp2/3 complex when ∼70 molecules of WASp are recruited to endocytic sites (Figures 6B and 6C, 8A(II) and 8B(II)) (Sun et al., 2017). These properties suggest that ∼70 WASp molecules are key for establishing a WASP-Arp2/3 complex stoichiometry sufficient for a burst of actin assembly (Case et al., 2019).

Second, the two yeasts recruit type I myosin and WASP to endocytic sites differently. The assembly/disassembly profiles of GFP-spWsp1 and spMyo1-GFP are similar in fission yeast (Figure 6B to 6E, Figure 8A), while scMyo3/5 appears at endocytic sites more than 10 s after scLas17 in budding yeast (Figure 8B(I)) (Sun et al., 2006).

Third, WASp-mediated actin assembly initiates membrane deformation in both yeasts. Endocytic membrane invagination is first detected when spWsp1 peaks and actin assembles to ∼50% of its maximum (Figure 6B and 6C, and Figure 8 A(III), 8B(III)).

Fourth, although both yeasts recruit WASp and myosin-I to endocytic sites, the yeast show different dependencies on the NPF activities of these two proteins. Budding yeast recruit a total of 300 molecules of two isoforms of myosin-I (vs. 170 in fission yeast) and the NPF activity of their CA motifs is important for robust internalization (Sun et al., 2006) but not in fission yeast (Fig7B and 7F). Nevertheless, complete deletion of fission yeast spMyo1 shows that, as in budding yeast, functions other than its NPF activity contribute to endocytosis (Arasada, 2011; Pedersen and Drubin, 2019; Sun et al., 2006). On the other hand, endocytic sites in fission yeast accumulate more WASp molecules (∼140 Wsp1) than in budding yeast (∼100 Las17) (Figure 2F), and the WASp NPF activity is crucial for endocytosis in fission yeast (Figure 7A, 7B and 7D) but not budding yeast (Sun et al., 2006).

Fifth, once membrane invagination begins, both WASp and myosin-I dissociate more quickly from actin patches in fission yeast (Figure 6C, Figure 6E and Figure 8(IV) and 8(V)) than budding yeast (Figure 8(IV) and 8(V)). However, the actin network peaks upon vesicle scission in both yeasts.

Finally, the two yeasts elongate the endocytic membrane invagination over the same time before scission, but the invagination grows two-fold faster and twice as long in fission yeast.

Our studies bring together two decades of research on endocytosis in yeast to illustrate how 300 million years of divergent evolution has resulted two systems using essentially the same complex set of conserved proteins somewhat differently to achieve a common result, formation of an endocytic vesicle. Further work will reveal the specific properties and advantages of each design solution and will compare the designs of these two systems with those in other organisms and different cell types in multicellular organisms.

## Experimental Procedures

### Media and Strains

*S. cerevisiae* and *S. pombe* strains were both grown in fission yeast standard rich media (YES 225, Sunrise Science Product) or synthetic media (EMM, MP Biomedicals).

### Preparation of 120-sfGFP-tagged nanocages from *E.coli*

10 O.D._600_ (optical density) *E. coli* expressing sfGFP-tagged nanocages were collected and washed twice with, and then diluted into, 500ul EMM media containing 50mM Tris (pH 7.4) and one protease inhibitor tablet (Roche). Sonication (1s on and 1s off for 1min, then on ice for 1min, repeating for 3 times) was used to break cells. The sonicated *E. coli* were centrifuged for 5 min at 14,000 rpm at 4°C and the supernatant was then collected. The supernatant was further diluted 10 times into the EMM media; we found that maintaining a pH in the range of 7.0 is important for this standard to accurately reflect intracellular molecule numbers. 5ul diluted supernatant was transferred onto a coverslip and the nanocages were allowed to settle for 10 min prior to imaging.

### Fluorescence microscopy

Fluorescence microscopy was performed using a Nikon Eclipse Ti microscope (Nikon Instruments, Melville, NY) controlled by Metamorph software (Molecular Devices, Sunnyvale, CA) and equipped with a Plan Apo VC 100×/1.4 Oil OFN25 DIC N2 objective (with Type NF immersion oil, Nikon), a Perfect Focus System (Nikon), and a Neo sCMOS camera (Andor Technology Ltd., South Windsor, CT) (65 nm effective pixel size). For live cell imaging, cells were grown overnight in YES 225 rich media. The saturated overnight cell culture was diluted into EMM synthetic media and cells were grown to early log phase at 25°C. When *S. pombe* and *S. cerevisiae* strains were imaged simultaneously, equal numbers of the two yeasts were mixed together just before imaging. The cells were transferred to a 1% agarose pad made from EMM synthetic media. All imaging was done at the equatorial plane of the cells at room temperature. For single-channel live-cell imaging, images were acquired continuously at 2-9 frames/s. Two-channel movies were made using the SPECTRA X Light Engine (Lumencor, Beaverton, OR) for excitation with a 524/628 nm dual-band bandpass filter for GFP/mCherry emission (Brightline, Semrock, Lake Forest, IL). Time to acquire one image pair was 137ms.

### Image analysis

Image J software was used for general processing of images and movies, such as background subtraction, photobleaching correction(Kaksonen et al., 2003). A median filter was used to subtract local background around patches as described in (Picco and Kaksonen, 2017). Particle Tracker plugin was used to track endocytic patch protein dynamics (Sbalzarini and Koumoutsakos, 2005) (Figure 1-figure supplement3). The trajectory selection was confirmed by visual inspection. Trajectory that any point in their lifetime were too close to another patch to be clearly resolved were excluded from our analysis. The sample size for each experiment is shown on the figure or/and in the figure legend. The sample sizes used in this study are more than what have been used in other landmark papers in this type of study (Kaksonen et al., 2003; Picco et al., 2015; Sirotkin et al., 2010). For each pair of variables, pooled data of analysis were compared by a two-sided Mann-Whitney test using the Prism 8 graphing software.

We built a custom data analysis pipeline (https://github.com/DrubinBarnes/YeastTrackAnalysis) that operates on individual tracks listed in the trajectory selection. For each track, the intensity values (I_t_), and the positions in x (x_t_) and y (y_t_) for each time point (t) of the track are recorded. Typical analysis pipelines for CME tracks [Berro and Pollard 2014, Picco et al. 2015] calculate track alignments by minimizing the least square difference between the time courses of track intensities (I_t_). This methodology makes the implicit assumption that the track intensity time course is regular from track to track over the whole duration of the measurement. However, different CME proteins might show different dynamics during their intensity time course and, depending on their role in CME, might show regularity patterns other than intensity profiles, such as protein displacement.

To identify such patterns, our pipeline is split into two parts: 1) In a first data preprocessing and examination step, multiple characteristics are quantified for each track (movement speed, time of 50% intensity, time of maximum intensity, etc.). This first step makes it possible to search for and find patterns and regularities among the recorded tracks. In step 2, alignments are calculated based on the new observed patterns from step 1. As a result, conclusions can be drawn from an aligned and averaged set of tracks. In this study, two characteristics proved to be most useful for our alignments: inflection point (see below) and 50% raw intensity.

#### Inflection point

Quantified coat protein tracks can usually be divided into two phases: An initial, non-motile phase (a) during which protein accumulates on the plasma membrane with little detectable movement and a second phase (b) in which the protein moves, likely reflecting the time when the plasma membrane deforms into an invagination (b_1_), and when the freed endocytic vesicle pinches off and moves into the cytoplasm (b_2_). We used the transition point between these two phases (a, b_1_), which we refer to as the inflection point t_inflection_, for CME track alignment. To calculate t_inflection_, first, the frame by frame displacement *d_t_* was calculated for all subsequent timepoints t and t+1 in a track: d_t+1_ = sqrt[(x_t+1_+1-x_t_)^2^ +(y_t+1_-y_t_)^2^]. The signature behavior is that *d_t_* is flat in phase (a) and then suddenly shows a slope change upon the transition to phase (b_1_). To identify t_inflection_, *d_t_* is first linearized using linear regression d_t,linear_ = linregress(d_t_) and then d_t,linear_ is subtracted from *d_t_*. t_inflection_ now is the minimum in the resulting function: t_inflection_ = argmin(d_t_ -d_t,linear_)

#### 50% raw intensity

The time point for 50% raw fluorescence intensity t_0.5,raw_ can be helpful to align the tracks for endocytic proteins with a short lifetimes and/or complex displacement patterns.

This value is found at Intensity(t_0.5,raw_) = 0.5 * max(Intensity(t)).

#### Implementation

The pipeline is implemented as a python software library with a Jupyter based user interface, which allows user-friendly and interactive access to its functionality. The pipeline is open source under a Berkeley Software Distribution (BSD) 3-clause license and can be found on Github: https://github.com/DrubinBarnes/YeastTrackAnalysis.

### 3D-STORM imaging and analysis

GFP nanobody Alexa Fluorphore 647 conjugation: A 100 µl GFP nanobody solution at 71µM (gt-250, ChromoTek) was conjugated to the Alexa Flurophore™ 647 NHS ester (A20006, Thermo Fisher Scientific) following standard procedures (Mund et al., 2014). After purification as described by (Mund et al., 2014) we obtained an AF647 GFP-nanobody solution with a concentration of 49.3 µM and a labeling ratio of approximately 1.2 dye per nanobody.

Fission yeast fixation and AF647 GFP-nanobody labeling: fission yeast cells were fixed and labeled following a method commonly used for good actin structure preservation (Kaplan and Ewers, 2015). Low speed centrifugation (1000 rpm) was used to pellet yeast cells. Paraformaldehyde was directly added to the culture to a 4% final concentration, followed by 15 min incubation. Cells were washed twice with CS (cytoskeleton) buffer (10mM MES, 150mM NaCl, 5mM EGTA, 5mM Glucose, 5mM MgCl_2_, 0.005% NaN3, pH6.1) containing 50 mM NH_4_Cl, for 5 mins with gentle shaking. The cells were collected and resuspended with CS buffer contain 5% Bovine serum albumin (BSA) and 0.25%(vol/vol) Triton X-100. After 30min gentle shaking, AF647 GFP-nanobody was added to the fixed cells to a final concentration of 0.5 µM. The fixed cells were incubated with AF647 GFP-nanobody with gentle shaking overnight at 4°C.

Alexa Fluorphore 647-labeled cell samples were mounted on gel pads with STORM imaging buffer consisting of 5% (w/v) glucose, 100 mM cysteamine, 0.8 mg/mL glucose oxidase, and 40 µg/mL catalase in 1M Tris-HCl (pH 7.5) (Huang et al., 2008; Rust et al., 2006). Coverslips were sealed using Cytoseal 60. STORM imaging was performed on a homebuilt setup based on a modified Nikon Eclipse Ti-U inverted fluorescence microscope using a Nikon CFI Plan Apo λ 100x oil immersion objective (NA 1.45). Dye molecules were photoswitched to the dark state and imaged using a 647-nm laser (MPB Communications, Quebec, Canada); these laser beam passed through an acousto-optic tunable filter and an optical fiber into the back focal plane of the microscope and onto the sample at intensities of ∼2 kW cm^-2^. A translation stage was used to shift the laser beams towards the edge of the objective so light reached the sample at incident angles slightly smaller than the critical angle of the glass-water interface. A 405-nm laser was used concurrently with the 647-nm laser to reactivate fluorophores into the emitting state. The power of the 405-nm laser was adjusted (typical range 0-1 W cm^-2^) during image acquisition so that at any given instant, only a small, optically resolvable fraction of the fluorophores in the sample were in the emitting state. For 3D STORM imaging, a cylindrical lens was inserted into the imaging path so that images of single molecules were elongated in opposite directions for molecules on the proximal and distal sides of the focal plane (Huang et al., 2008; Rust et al., 2006). The raw STORM data was analyzed according to previously described methods(Huang et al., 2008; Rust et al., 2006). Data were collected at a frame rate of 110 Hz, for a total of ∼60,000 frames per image.

Endocytic sites were first identified through signals in the 488nm channel. Conventional images of yeast containing puncta or bright spots were then overlaid with the image in the super-resolution channel by adding the super-resolution image to the conventional image. If signals in both the conventional and super-resolution channel overlapped, the structure in the super-resolution channel was cropped and the corresponding molecule list (includes X, Y, Z positions of all molecules in the reconstructed image) for further data analysis. If no signal overlap was observed between the conventional and super-resolution channel, the site was not subjected to analysis. The puncta in crowded regions of the cell could not be clearly resolved as a single endocytic site and therefore were not included in our analysis.

The single molecule positions for the structures of interest obtained from the reconstructed STORM image were loaded into MATLAB. A rotation matrix was applied to the molecule position list such that the invagination site was rotated to a horizonal orientation. The length of the invagination was measured by computing the distance between every pair of molecules within a set threshold (∼0.01 difference in pixel size) along the y-axis, until the longest distance was found. The distance between the two molecules was calculated by the distance formula (d = √ ((x_2_-x_1_)^2^ + (y_2_-y_2_)^2^)). The length, in pixel size, was then converted to nanometers using the conversion scale 1 pixel = 160nm. The position of the plasma membrane relative to the endocytic structures was estimated by the background signals showing the cell shape.

## Acknowledgements

We thank members of the Drubin/Barnes laboratory for helpful discussions. We greatly appreciate Dr. Jonathan Wong and Dr. Meiyan Jin for critically reading our manuscript. We are grateful to Paula Real Calderon in Fred Chang’s laboratory for advice on fission yeast molecular biology. We also thank Dr. Samuel J Kenney for help in establishing the 3D-STORM experimental setup. We also thank Dr. Andrea Picco for his advice on particle tracking analysis. This work was supported by NIH grant R35GM118149 to D.G.D. Dr. Tom Pollard was a Miller Visiting Professor at UC Berkeley.

## Supplemental Figure Legends

**Figure 1-table supplement 1.** Comparison of peak protein numbers in actin patches.

| Protein | S. pombe |  |  | S. cerevisiae |  |  |
| --- | --- | --- | --- | --- | --- | --- |
|  | Pollard lab: Sirotkin (2010)<sup>a</sup>; Arasada (2011)<sup>b</sup>; Chen (2013)<sup>c</sup>; Epstein (2018)<sup>d</sup> | Basu (2011) | Sun<br>This study | Galletta (2012) | Picco (2015) | Sun<br>This study |
| Actin | 7500<sup>a</sup>; 4050<sup>b</sup> |  |  |  |  |  |
| Arp2/3 complex subunits | 320, 320, 320<sup>a</sup> | ~150 | **304**<br>**(302)** | 300 | 210 | **294** |
| WASp/Las17 | 230<sup>a</sup>; 140<sup>b</sup>; 135<sup>c</sup> | 125 | **138** |  | 45 | **102** |
| WIP/Vrp1 | 140<sup>a</sup> |  | **95** |  |  | **78** |
| Myosin-I/<br>Myo3-5 | 400<sup>a</sup>; 170<sup>b</sup>; 240<sup>c</sup> |  | **170** |  | Myo5: 125 | **Myo5: 200**<br>**Myo3: 100** |
| Fimbrin/Sac6 | 910<sup>a</sup>; 800<sup>c</sup>; 1000, 1600<sup>d</sup> |  | **675** |  |  | **545** |
| HIP1R<br>End4/Sla2 | 160<sup>a</sup>; 70<sup>c</sup> |  | **124** |  | 30, 40 | **133** |
| Intersectin<br>Pan1/End3 | Pan1: 260<sup>a</sup>; 160<sup>c</sup> |  | **Pan1: 219** |  | End3: 55 | **End3: 100**<br>**Pan1: 131** |
| Sla1 |  |  |  |  | 85 | **168** |
| ABP1 |  |  |  | 810 | 423 | **800** |
| Clathrin HC | 40<sup>a</sup> |  |  |  |  |  |
| Clathrin LC | 40, 30<sup>a</sup> | ~30-40 |  |  |  |  |
| Capping protein | 230<sup>a</sup> |  |  | 200 |  |  |
| App1 | 150<sup>a</sup> |  |  |  |  |  |
| Coronin | 490<sup>a</sup>; 320<sup>b</sup> |  |  |  |  |  |
| Twinfillin | 210<sup>a</sup> |  |  |  |  |  |
| F-BAR Cdc15 | 125<sup>b</sup> |  |  |  |  |  |
| F-BAR Bzz1 | 90<sup>b</sup> |  |  |  |  |  |
| Dip1 |  | 20 |  |  |  |  |

**Calibration methods:** Sirotkin, Arasada, Chen, Epstein: a range of internal standards tagged with YFP or GFP; Basu: comparison with actin patches in other cells marked with Arp2/3 complex subunit mYFP-arc5 assumed to be 330 molecules per patch from Sirotkin et al. (2010); Sun: External standard nanoparticle with 120 eGFPs; Galletta: internal Cse4 standard; Picco: internal Nuf2 standard which was calibrated as a ratio with Cse4.

**Figure 1-figure supplement 1.**
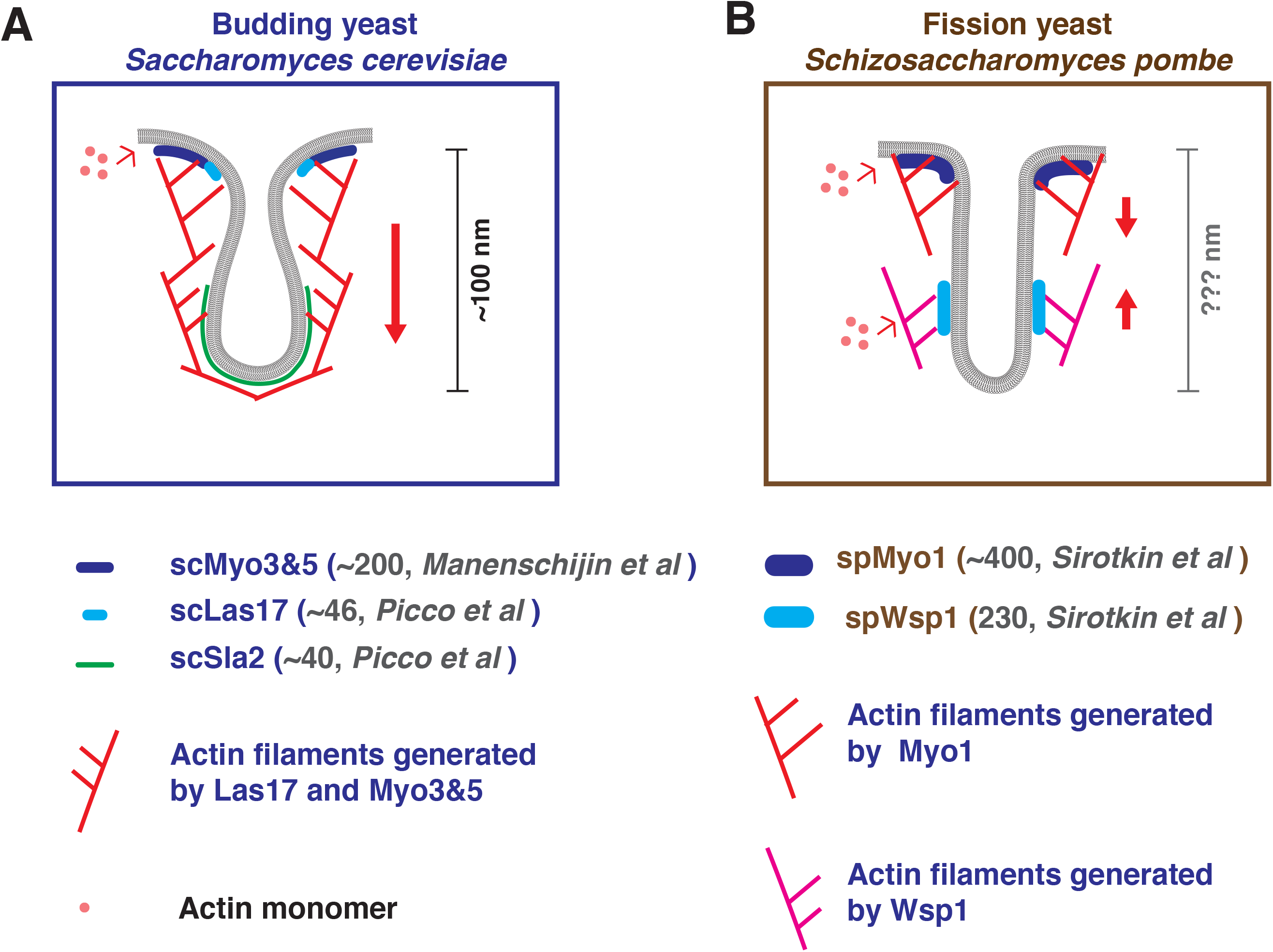
Endocytic sites in budding and fission yeast: A brief summary of what has been reported previously. The endocytic actin machinery on invaginated membranes in budding (A) or fission (B) yeast. A, In budding yeast, actin filaments are proposed to be nucleated near the base of the invagination. Continued polymerization driven by the nucleation promoting activity (NPF) of scMyo3/5 and scLas17 pushes the actin network toward cytoplasm (Sun et al., 2006). The endocytic membrane is coupled to the actin network by coat proteins such as Sla2 and is pulled inward with the actin network (Kaksonen et al., 2003; Sun et al., 2005). The membrane invagination is ∼100nm deep at the time of vesicle scission (Kukulski et al., 2012). B, In fission yeast, a two-zone model proposes that actin nucleation by spMyo1 and spWsp1generate two independent actin networks that push against each other and pull the tip of the invagination into the cytoplasm(Arasada and Pollard, 2011). The depth of the fission yeast membrane invagination is not known. Note, the previously measured absolute molecular numbers of indicated protein homologues differ greatly between the two yeasts (Picco et al., 2015; Sirotkin et al., 2010).

**Figure 1-figure supplement 2.**
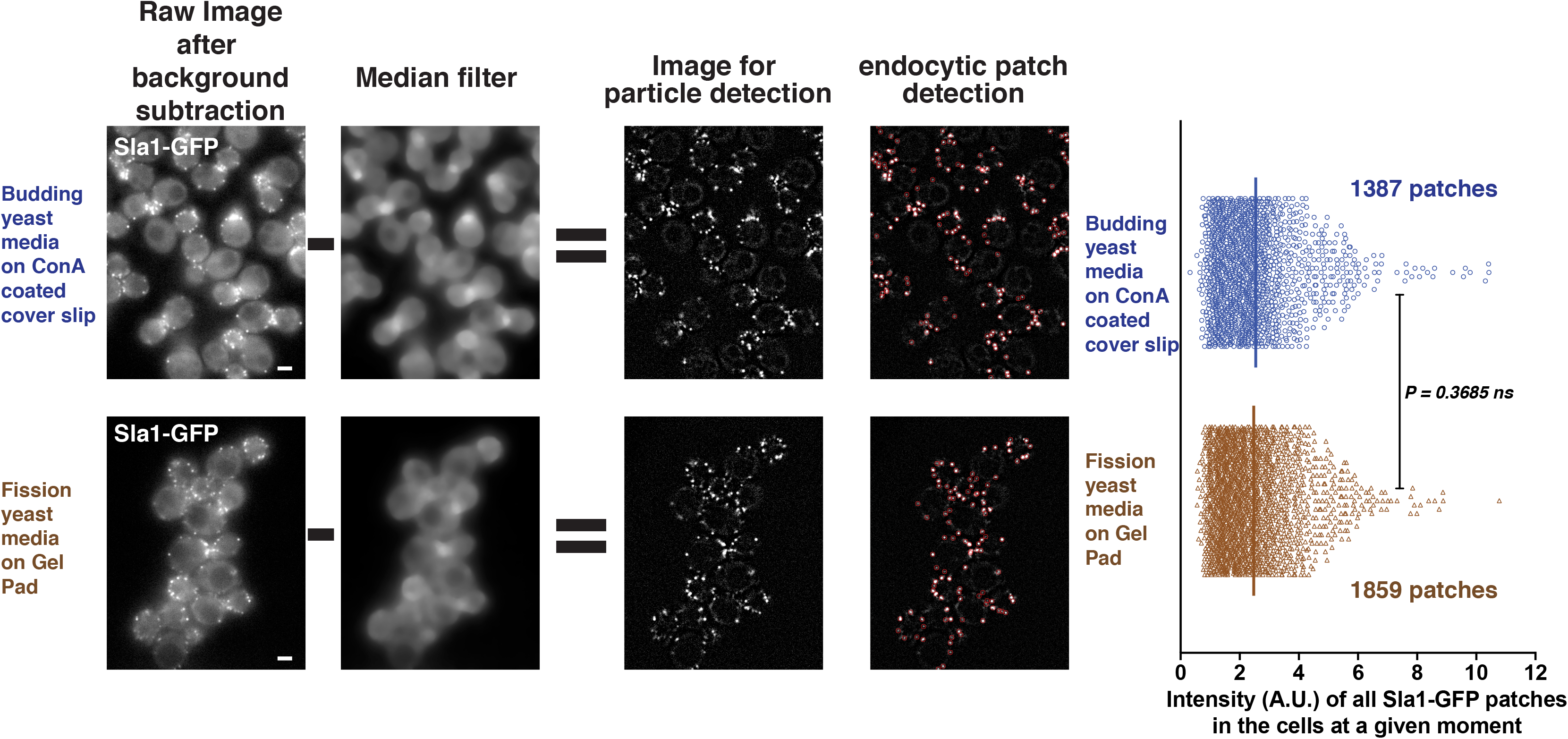
Quantitative comparison of fluorescently-tagged scSla1 in budding yeast cultured in different media. Budding yeast expressing *SLA1-GFP* were cultured in budding yeast (upper panel) or fission yeast (lower panel) media. The cells were then imaged by the methods commonly used for budding or fission yeast, respectively. The images were processed and analyzed using the Particle Tracker plugin in ImageJ software (see methods). To precisely compare the number of endocytic proteins at endocytic sites under these different growth conditions, local background correction by median filter subtraction (Picco and Kaksonen, 2017) was applied to the live cell images after general background subtraction and photobleaching correction. Red circles in right panels indicate the endocytic events automatically detected by the tracking program. The fluorescence intensities of all the detected events were measured and compared. Scale bars are 2µm.

**Figure 1-figure supplement 3.**
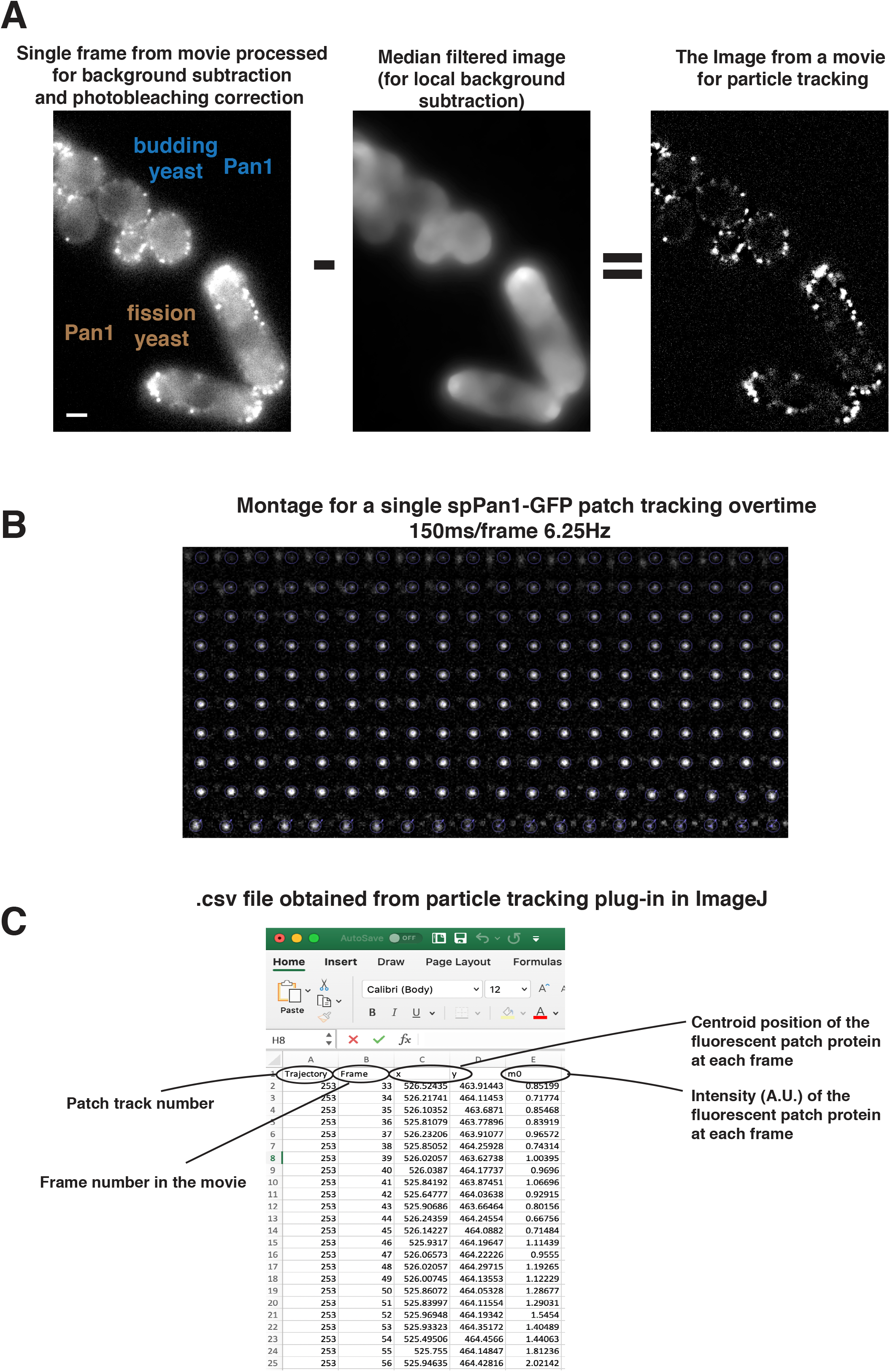
Live cell imaging data processing and analysis by Particle Tracker plugin. A, Single frame from movie processed for background subtraction and photobleaching correction (left panel). A median filter was used to compute the local background surrounding the endocytic patches (middle panel). The median filtered image was subtracted from the processed image, resulting the image (right panel) for particle tracking. B, Time series showing intensity and movement of a single endocytic event over its full life-time using Particle Tracker. C, The .csv file generated by particle tracking analysis. Scale bars are 2µm.

**Figure 2-figure supplement 1.**
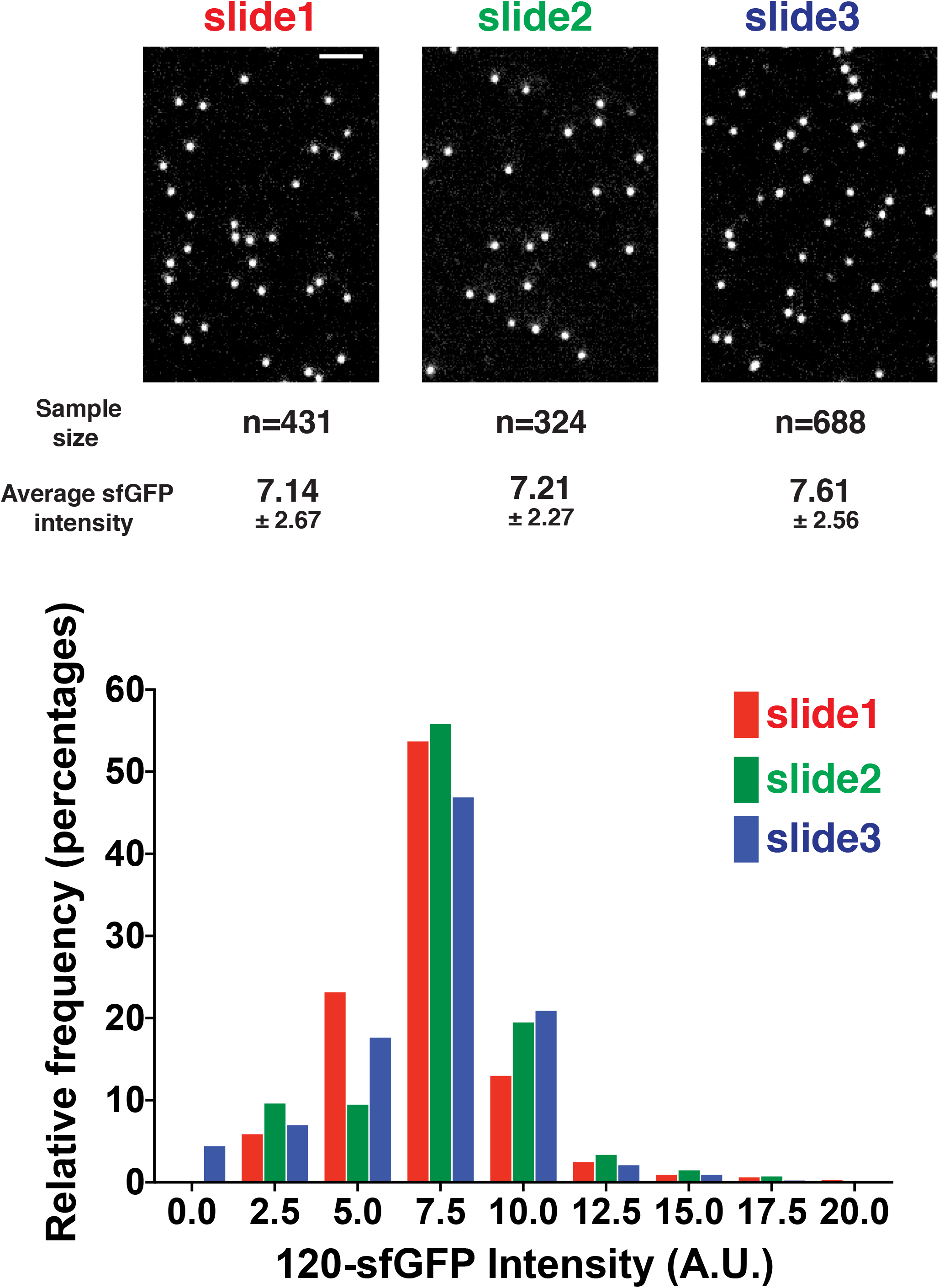
Quantitative comparison of 120-sfGFP-tagged nanocages prepared on different days. 120-sfGFP-tagged nanocages were prepared and imaged in three different days using the same method and conditions. The fluorescence intensities of the 120-sfGFP-tagged nanocages were measured and compared. Scale bars are 2µm.

**Figure 2-figure supplement 2.**
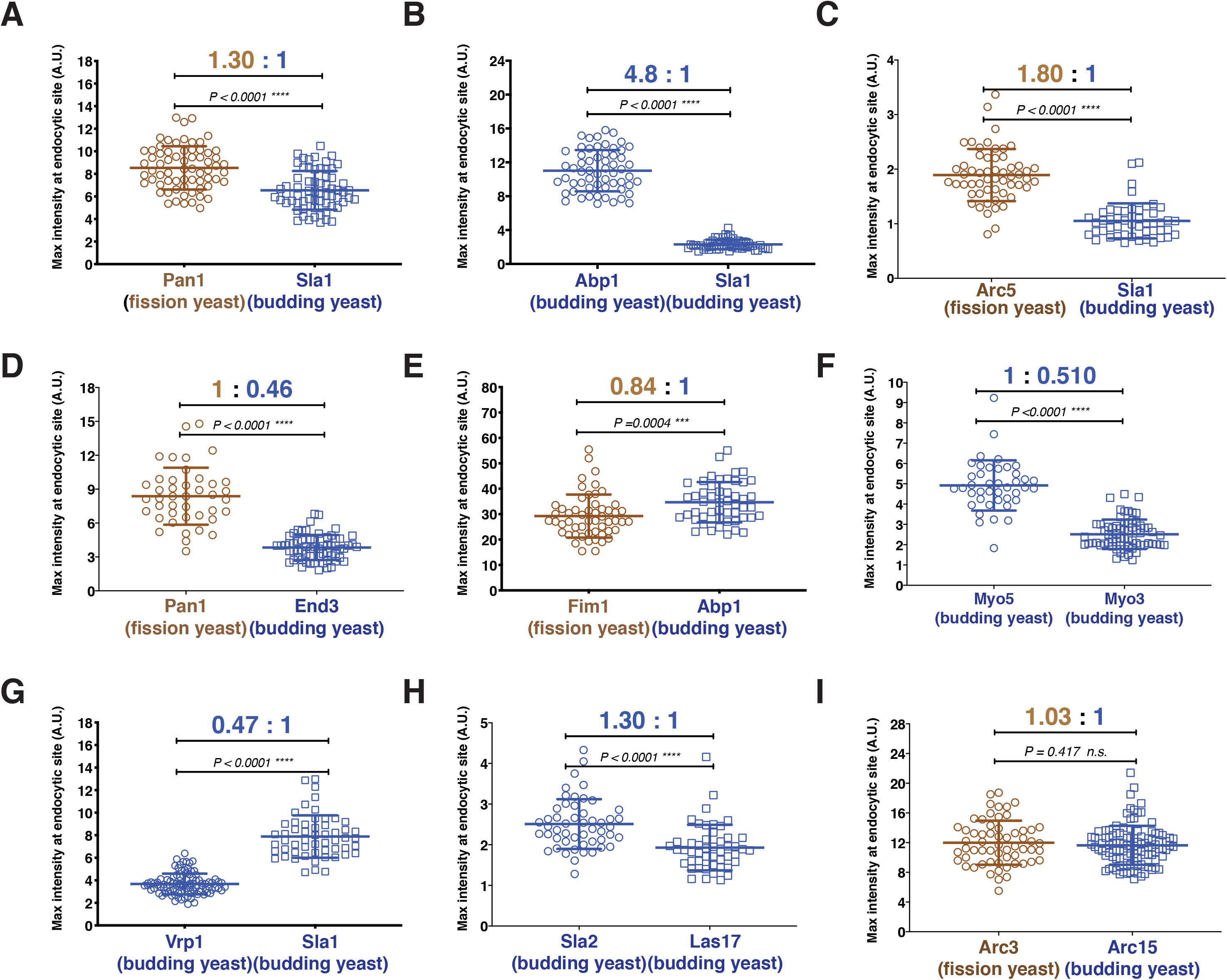
Quantitative comparison of the maximum number of homologous proteins at endocytic sites in budding and fission yeast. A-I, Cells expressing either indicated proteins were mixed and then imaged over time. The resulting movies were analyzed using a particle tracking program. The maximum fluorescence intensities was measured and compared for the indicated protein pairs.

**Figure 2-figure supplement 3.**
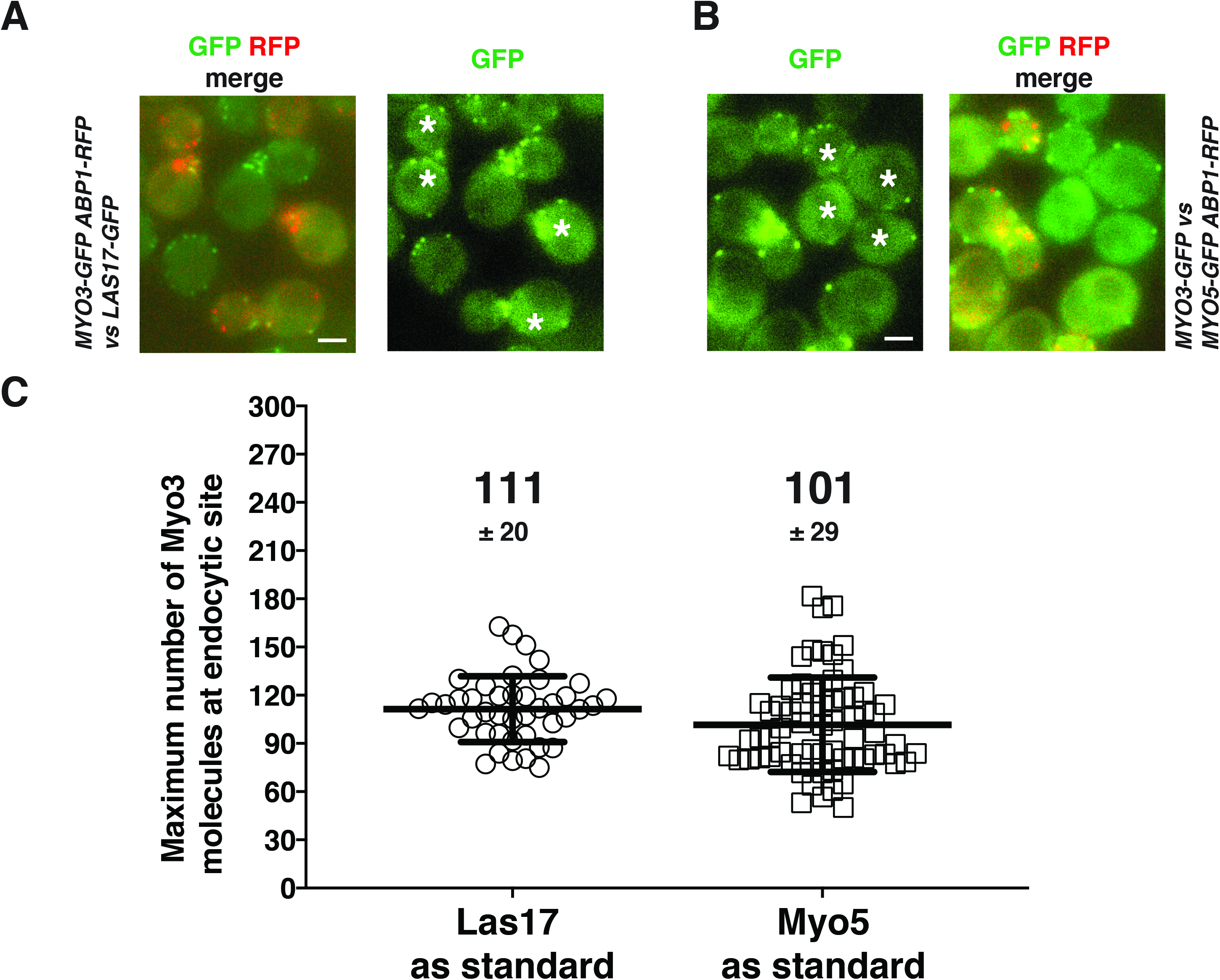
Determining the maximum number of Myo3-GFP molecules using Sla1-GFP and Las17-GFP as standards. A, Determining the maximum number of Myo3-GFP molecules using Las17-GFP as a standard. Two-color image of *MYO3-GFP ABP1-RFP* cells and *LAS17-GFP* cells (left panel). Single frame of a movie of *MYO3-GFP ABP1-RFP* cells and *LAS17-GFP* cells simultaneously imaged in the GFP channel (right panel). The dynamics of GFP-labeled protein patches were analyzed using the Particle Tracker plugin in ImageJ software and the maximum fluorescence intensity was measured and compared between the indicated proteins (C). B, Determining the maximum molecular numbers of Myo3-GFP using Myo5-GFP molecules for their subsequent use as standards. Two-color image of *MYO5-GFP ABP1-RFP* cells and *MYO3-GFP* cells (left panel). Single frame from a movie of *MYO5-GFP ABP1-RFP* and *MYO3-GFP* cells imaged simultaneously in the GFP channel (right panel). The dynamics of GFP-labeled protein patches were analyzed using the Particle Tracker plugin in ImageJ software and the maximum fluorescence intensity was measured and compared between the indicated proteins (C). C, The molecular numbers were calculated by ratiometric fluorescence intensity comparison using data acquired in A and B. The asterisks represent *LAS17-GFP* cells in (A) and represent *MYO3-GFP* cells in (B). Scale bars are 2µm.

**Figure 3-figure supplement 1.**
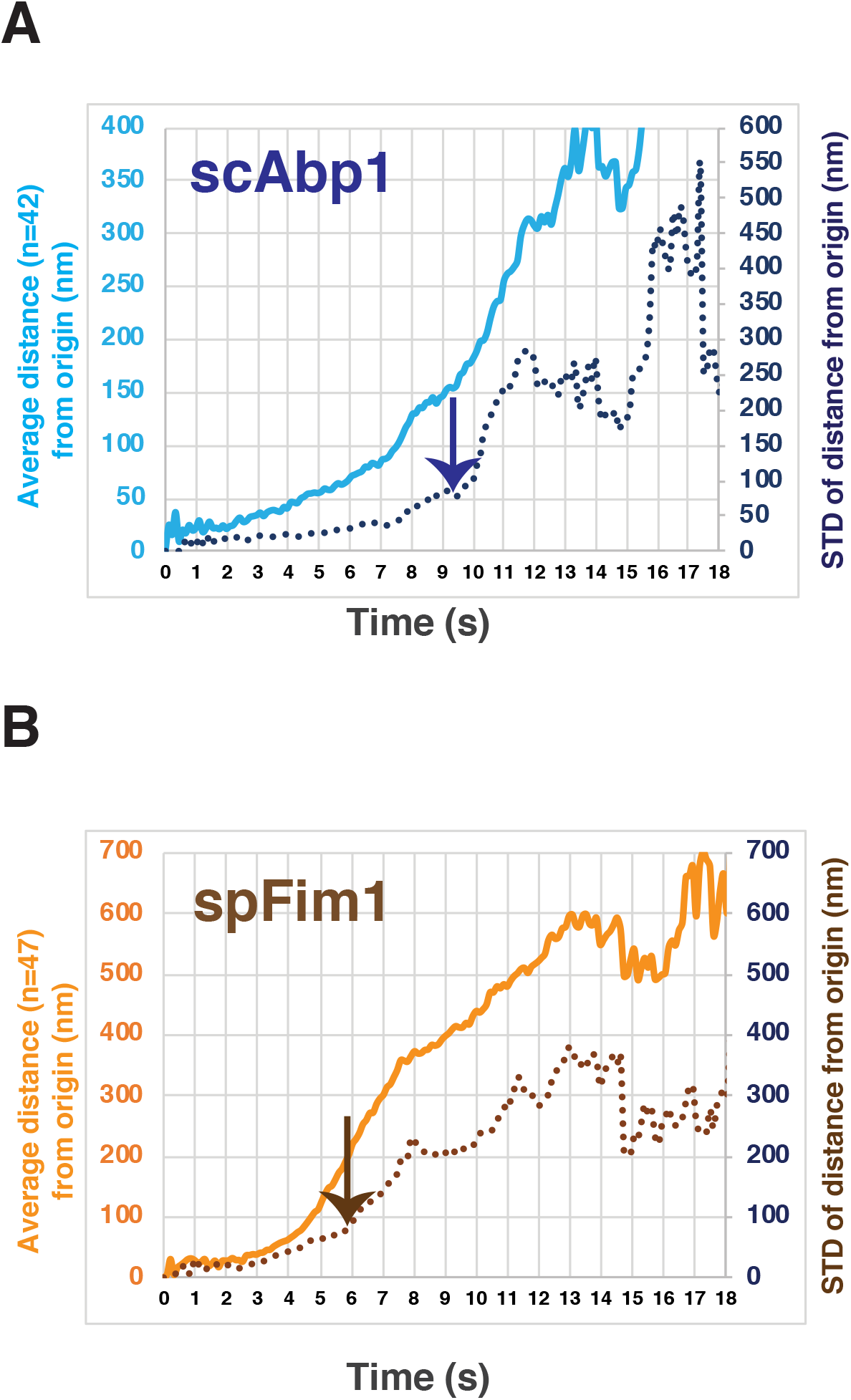
Predicting the timing of endocytic vesicle scission by the changes of the standard deviation of actin patch displacement. A, The standard deviation (STD) of actin patch displacement starts to greatly increase when scAbp1 patches move ∼125nm away from their origin. The light blue line represents the average displacement from the origin. The dark blue dotted line represents the STD of displacement from the origin. The arrow indicates the timing of the predicted scission event. B, The STD of actin patch displacement starts to greatly increase when scFim1 patches move ∼200nm away from their origin. The light brown line represents the average displacement from the origin. The dark brown dotted line represents the STD of displacement from the origin. The arrow indicates the timing of the predicted scission event.

**Figure 5-figure supplement 1.**
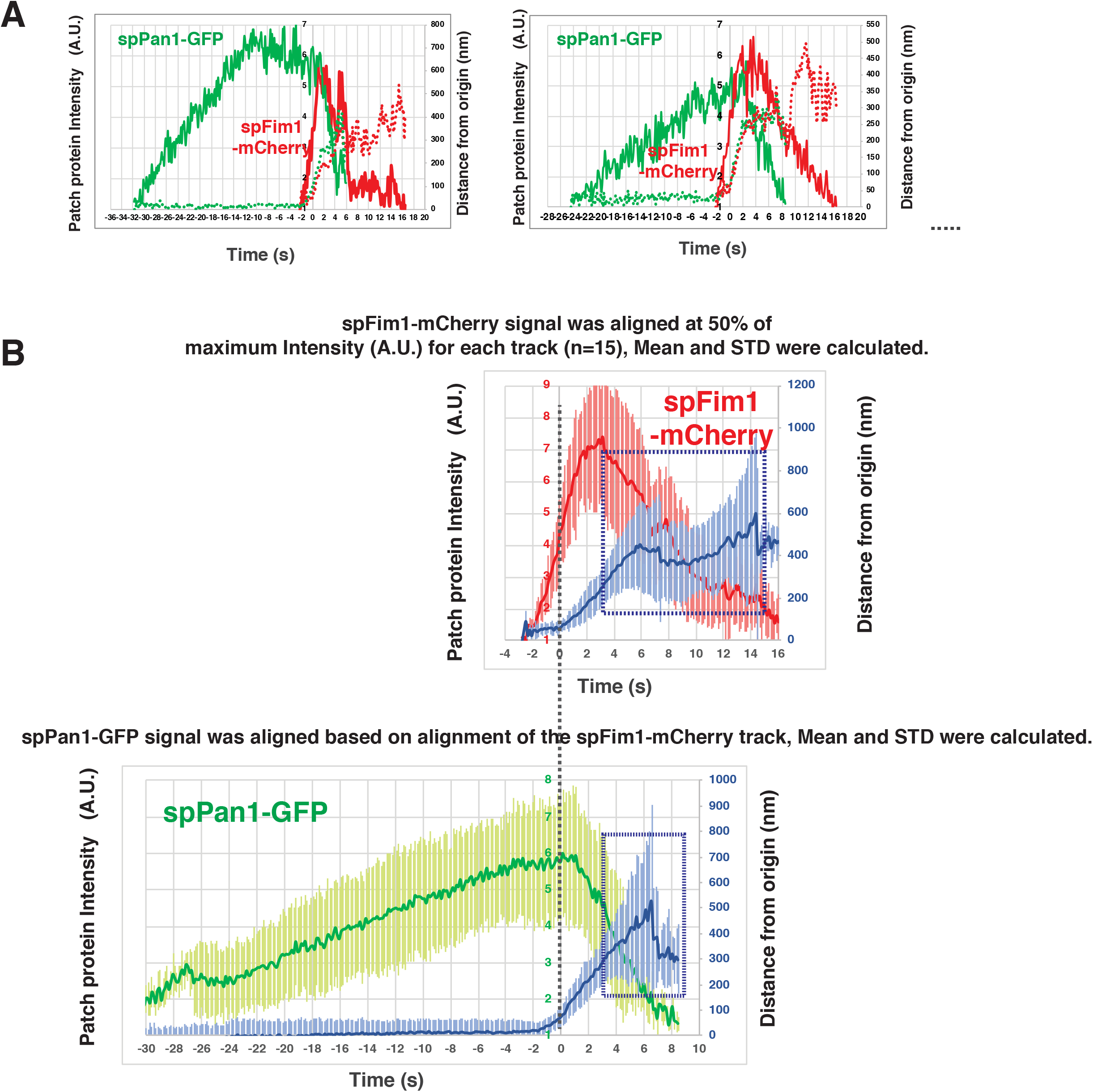
Alignment and quantification of average trajectories for spFim1-mCherry and GFP-spPan1 in fission yeast. A, Alignment of intensity and displacement of spPan1-GFP and spFim1-mCherry for a single endocytic event. Two examples are presented. B, Alignment of average intensity and displacement plots for spFim1-mCherry (upper graph) and GFP-spPan1 (lower graph) patches. Dotted vertical line aligns two graphs. Dotted boxes indicate inferred movement after scission.

**Figure 5-figure supplement 2.**
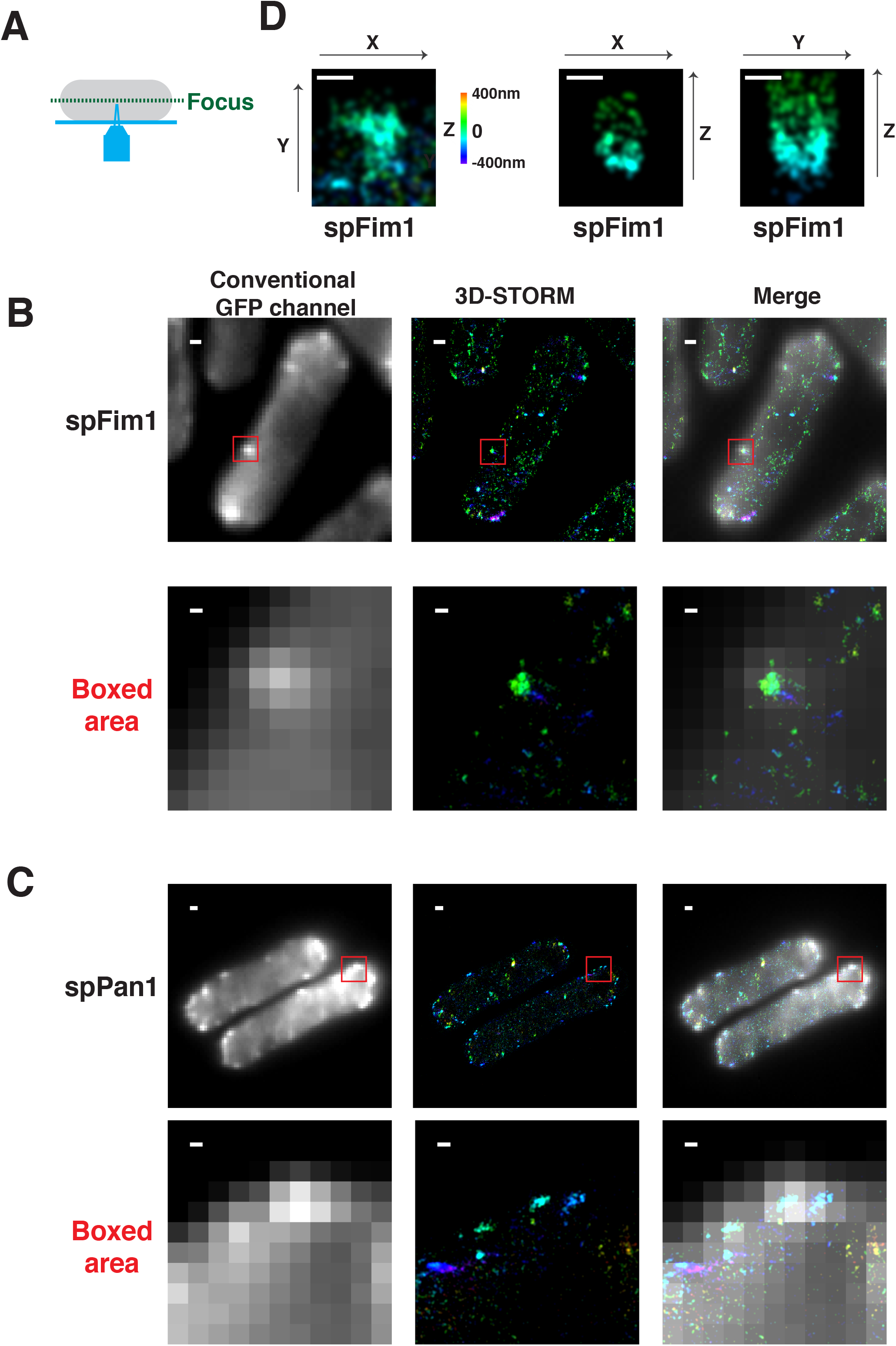
A, Chemically fixed fission yeast cells expressing GFP-tagged proteins were labeled with AF647-conjugated anti-GFP nanobodies and imaged at the equatorial plane. B, Conventional and 3D-STORM image for fixed fission yeast cell expressing spFim1-GFP and labeled using AF647-conjugated anti-GFP nanobodies. C, Conventional and 3D-STORM image of fixed fission yeast cell expressing spPan1-GFP and labeled using AF647-conjugated anti-GFP nanobodies. D, 3D-STORM image of spFim1-GFP at an individual endocytic site. Images represent XY, XZ, or YZ dimensions. Note, the Z-axis information is represented by rainbow color. The scale bars on the images of whole cells are 1µm. The other scale bars in the boxed areas and D are 100nm.

**Figure 6-figure supplement 1.**
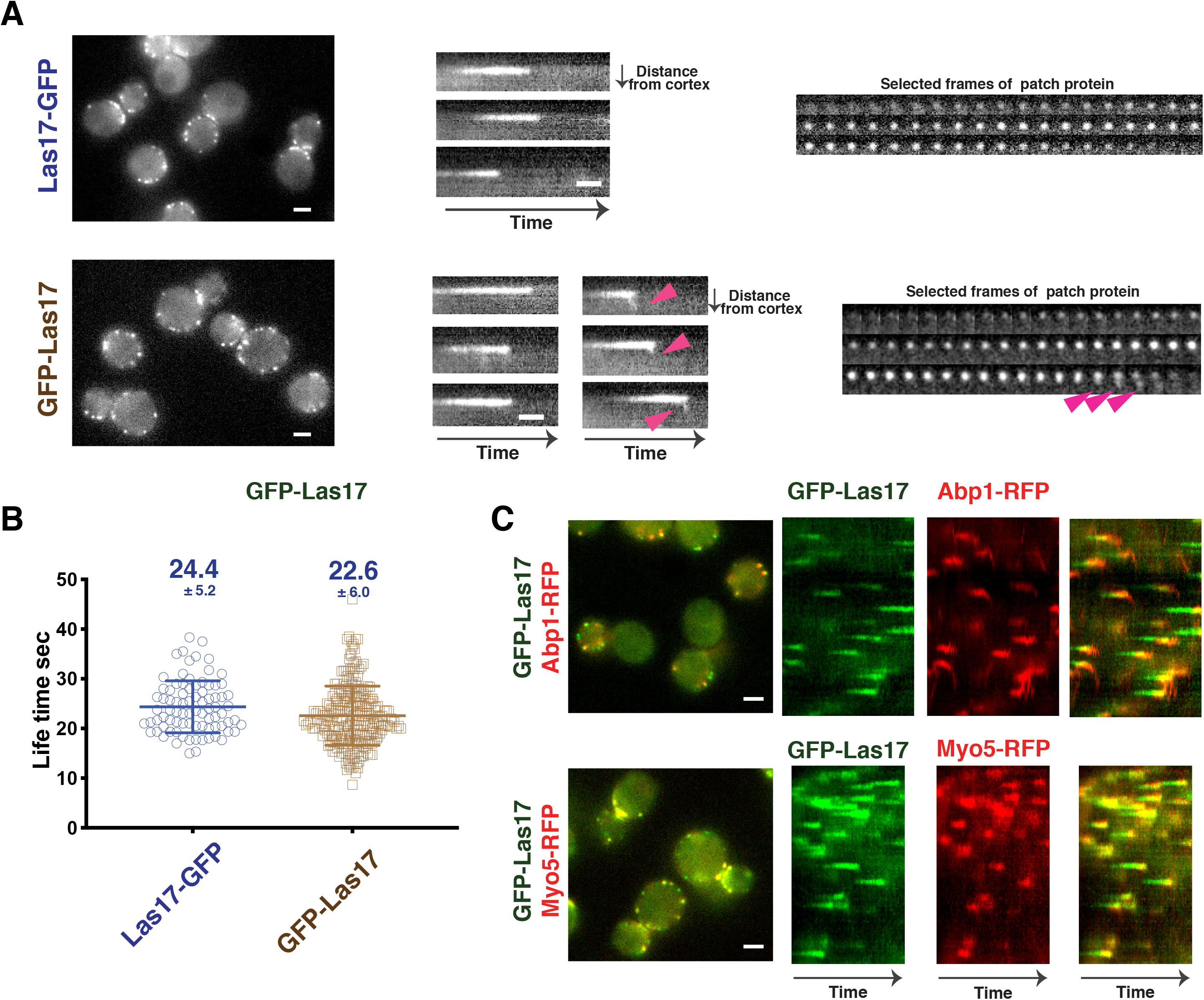
Dynamics of GFP-scLas17 and scLas17-GFP in budding yeast. A, Dynamics of scLas17-GFP and GFP-scLas17. Single frames (left) from movies, radial kymograph representations (Sun et al., 2017) (middle), and time series showing progression of a single endocytic event (right) for scLas17-GFP (upper panel) and GFP-scLas17 (lower panel). The arrow heads indicate that a small amount of GFP-scLas17 signal splits from the majority of the signal at the end of its life time. B, Life times of scLas17-GFP and GFP-scLas17 at endocytic sites. C, Single frames (left) from movies and circumferential kymograph representations (Sun et al., 2017) of *GFP-LAS17 ABP1-RFP* cells (upper panel) and *GFP-LAS17 MYO5-RFP* cells (lower panel). Scale bars are 2µm. Scale bars on kymograph are 10s.

**Figure 6-figure supplement 2.**
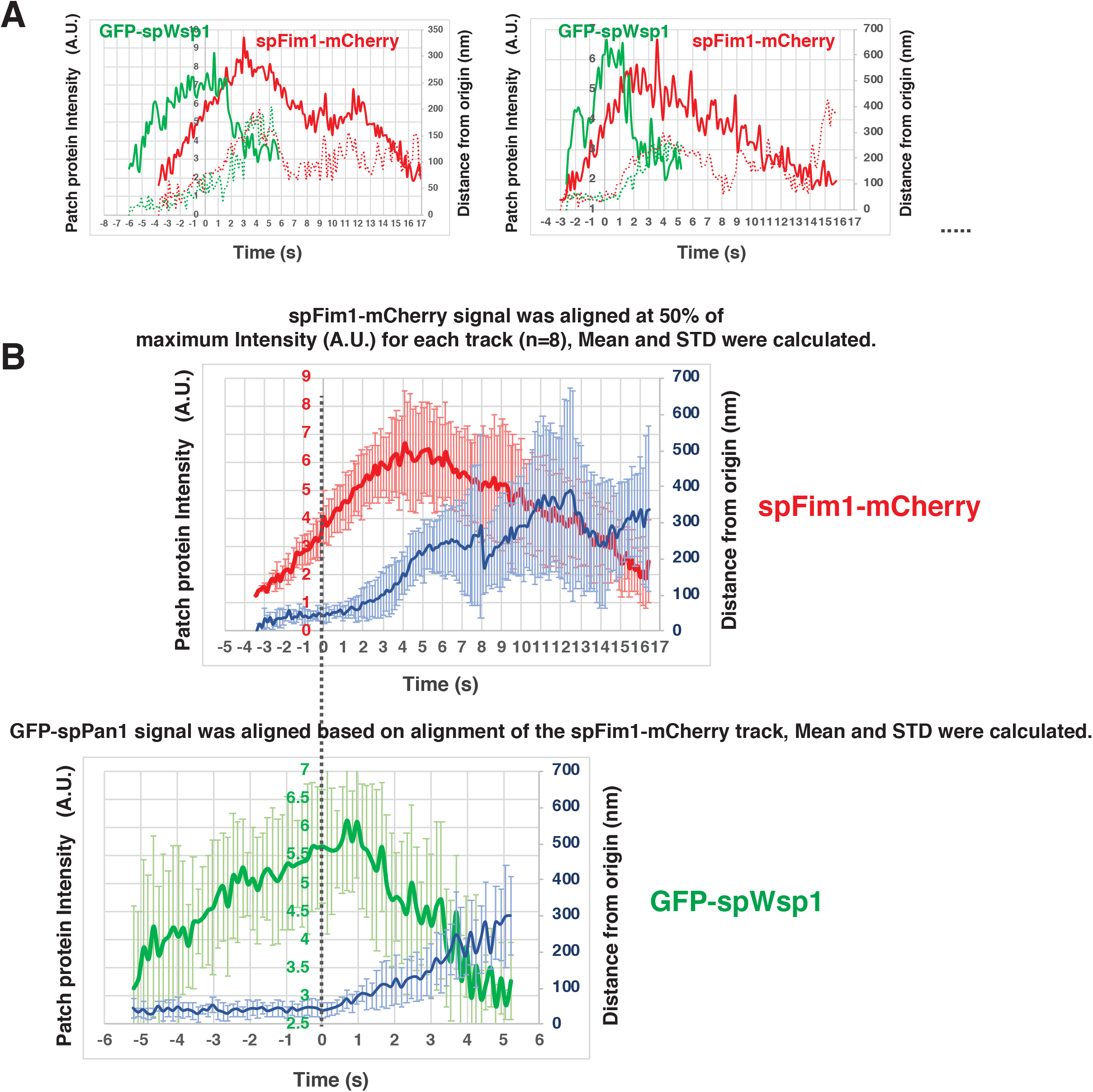
Alignment and quantification for average trajectories for spFim1-mCherry and GFP-spWsp1 in fission yeast. A, Alignment of intensity and displacement for GFP-spWsp1 and spFim1-mCherry for a single endocytic event. Two examples are presented. B, Alignment of average intensity and displacement of GFP-spWsp1 and spFIM1-mCherry patches.

**Figure 6-figure supplement 3.**
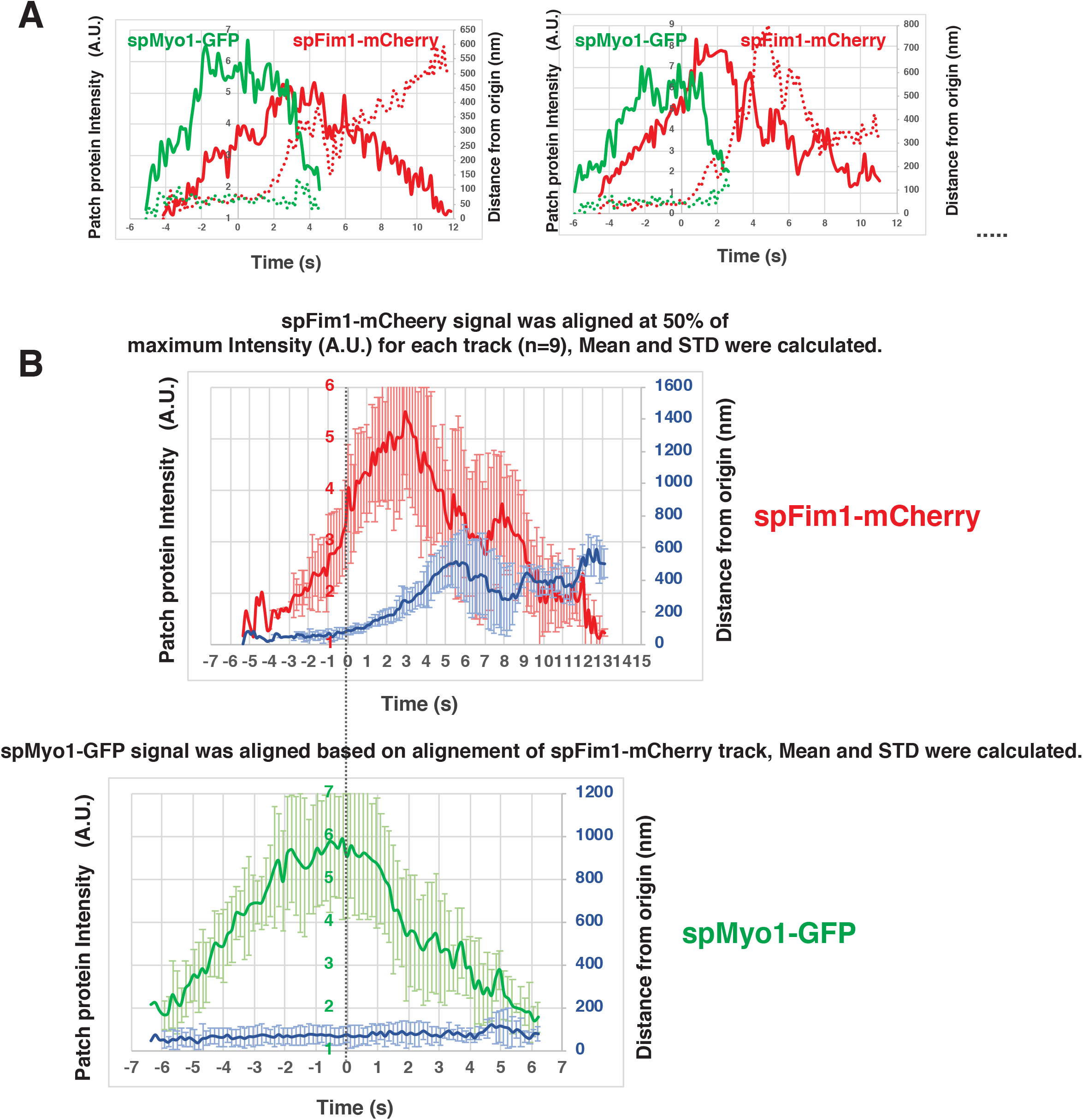
Alignment and quantification for average trajectories for spFim1-mCherry and spMyo1-GFP in fission yeast. A, Alignment of intensity and displacement for spMyo1-GFP and spFim1-mCherry for a single endocytic event. Two examples are presented. B, Alignment of average intensity and displacement for spMyo1-GFP and spFIM1-mCherry patches.

## Supplemental Movie Legends

**Video 1,**

Dynamics of spWsp1-GFP and spFim1-mcherry in *spWSP1-GFP spFim1-mcherry S. pombe* cells. Time to acquire one image pair was 136 ms. Interval between frames is 548 ms.

**Video 2,**

Dynamics of spwsp1CAΔ-GFP and spFim1-mcherry in *spwsp1-CAΔ-GFP spFim1-mcherry S. pombe* cells. Time to acquire one image pair was 400 ms. Interval between frames is 460 ms.

**Video 3,**

Dynamics of spMyo1-GFP and spFim1-mcherry in *spMyo1-GFP spFim1-mcherry S. pombe* cells. Time to acquire one image pair was 200 ms. Interval between frames is 221 ms.

**Video 4,**

Dynamics of spMyo1CAΔ-GFP and spFim1-mcherry in *spmyo1CAΔ-GFP spFim1-mcherry S. pombe* cells. Time to acquire one image pair was 200 ms. Interval between frames is 221 ms.

